# Sperm hyperactivation drives a circling-and-wandering migration strategy

**DOI:** 10.1101/2025.07.03.662809

**Authors:** Meisam Zaferani, Yanis Baouche, Yamilka Lago-Alvarez, Anish Pandya, Soon Hon Cheong, Sabine Petry, Christina Kurzthaler, Howard A. Stone

## Abstract

During migration through the female reproductive tract, sperm undergo physiological changes known as capacitation, including a motility transition termed hyperactivation. Hyperactivation is essential for various aspects of fertilization, particularly effective migration within the tract. However, how hyperactivation facilitates this migration remains elusive. Here, we profiled bull sperm hyperactivation in Newtonian and complex fluids near microfluidic surfaces, mimicking generic swimming conditions in the tract. We identified three swim gaits: wandering (persistent random walks), circling, and an intriguing circling-and- wandering mode marked by stochastic transitions between the two. All gaits exhibit diffusive behavior over long time scales, with wandering showing a tenfold higher diffusivity than circling, and the diffusivity of circling-and-wandering falling in between. We found that while wandering sperm scatter from convex and concave surfaces, circling sperm become trapped around pillars, highlighting the distinctive nature of each phase. Additionally, stochastic simulations of swimming in porous media showed that as the geometrical complexity of the environment increases, circling-and-wandering outperforms either strategy alone in spreading through the media. Our findings suggest that while wandering promotes exploration and circling supports local exploitation, circling-and-wandering combines the strengths of both strategies by balancing exploration and exploitation to adapt motility, enhance migration, and potentially improve target search.

---

For fertilization, mammalian sperm navigate the complex environment of the female reproductive tract (FRT) and withstand selective pressures. This migration is regulated by cues provided by the tract. Previous studies have shown that when swimming progressively in straight lines, sperm migrate upstream in response to external fluid flows^1–3^. Furthermore, physical boundaries, such as the walls of microfluidic channels, direct the progressive motion of the sperm^4–7^. These observations suggest that the mucus flow and the architecture of the tract may modulate sperm migration, allowing them to traverse the lower regions of the tract and reach its upper part.

During progressive swimming, sperm motility consists of a slightly asymmetric two-dimensional (2D) flagellar beating pattern^8–11^, leading to helical motion in bulk fluid and circular motion near surfaces. Furthermore, sperm have been observed to roll unidirectionally around their longitudinal axis^12^. Previous observations on rodent sperm and computational analyses suggest that the slightly tilted head, with respect to the flagellar beating plane, causes this rolling motion^10, 13–16^.

However, as sperm enter the upper part of the tract, they become capacitated^17, 18^ and their flagellar beating patterns become highly asymmetric^19, 20^, shifting progressive motility to hyperactivated motility, which is characterized qualitatively by diverse movement patterns that trace erratic trajectories^18, 21–23^. It is known that hyperactivation is caused primarily by the activation of the CatSper channel, which is distributed throughout the flagellum^24–26^. The structure and function of this channel^27–29^, its evolutionary origin^30^, and its necessity for fertilization^31, 32^ are well established. In addition, hyperactivation – induced by chemical^33, 34^ and/or thermal^35, 36^ stimuli – is known to facilitate sperm migration within the geometrically complex upper FRT, lined with heterogeneous mucus^23, 26, 37–43^. However, a physical framework to elucidate how hyperactivation modulates sperm migration through rheological and geometrical complexities has yet to be developed.

Here, we show that hyperactivated motility manifests itself in distinct forms of near-surface motion depending on the rheological properties of the fluid. Combining microfluidic experiments, theoretical analysis, and stochastic simulations, we further explore the implications of these sperm motility patterns for migration within the FRT. This work refines the qualitative understanding of hyperactivation and lays the foundation for a physical model of mammalian sperm chemotaxis, a question that, unlike the well-studied chemotaxis in marine invertebrates^44–47^, remains open.

## Statistical profiling of hyperactivated motility

We characterized the motility of bull sperm in an otherwise quiescent fluid in a circular microfluidic chamber (diameter = 1200 *µ*m and height = 30 *µ*m), designed to mimic the high-aspect-ratio geometry of the sperm passage in the FRT, such as microgrooves and narrow spaces between mucosal folds^41, 48^. To induce sperm hyperactivation, 3.0 and 6.0 mM of caffeine, an established hyperactivation agonist^22, 49, 50^, were added to the medium. We used Tyrode’s albumin lactate pyruvate (TALP) as a standard Newtonian buffer, and to mimic the shear-thinning viscoelastic nature of mucus, we supplemented TALP with 1% polyacrylamide (PAM)^51^. Sperm motion was then recorded at 15 frames per second (fps) for durations of 3 to 15 minutes.

### Progressive motion in a Newtonian fluid

Bull sperm consist of a paddle-shaped head (of length *l*_*h*_ = 5 *µm*) and a flagellum whose wave-like bending induces swimming. This form of motility in TALP resulted in persistent movement on straight trajectories (referred to as ‘progressive’), near the top and bottom surfaces of the chamber, at a translational speed of 101 ± 11 *µ*ms^−1^. Their swimming was subject to small noise, with a rotational diffusivity of *D*_rot_ = 0.05 ± 0.02 s^−1^. This noise arises from active swimming, rather than from typical Brownian motion. Observing at high temporal resolution (100 fps) and using the paddle-shaped head, we identified a nearly two-dimensional flagellar beating pattern (Fig. S1**a**). We also measured the rolling frequency^16, 52^ as 9.0 ± 2.0 Hz (Fig. 1**a** and **b**, Movie S1). The rolling is presumably induced by the tilt of the sperm head relative to the flagellar beating plane, at an angle of *φ* = 13.0^°^ ± 3.5^°^ (Fig. S1**b**-**c**), which is consistent with the torque-free condition of microswimmers in viscous flows^13^.

**Figure 1.**
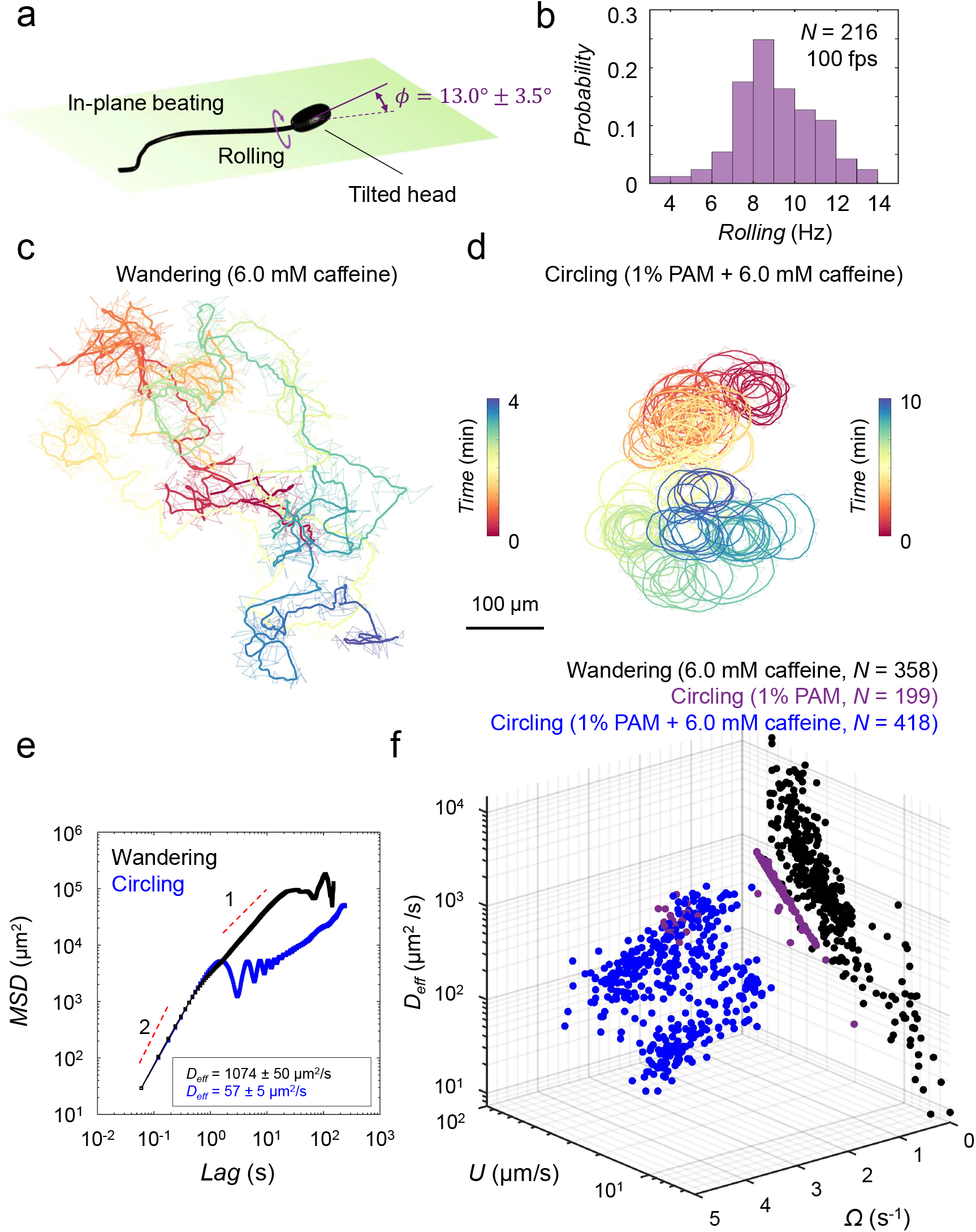
Statistical profiling of hyperactivated motility. (a) Schematic of bull sperm morphology and motility. The sperm head is slightly tilted out of the flagellar beating plane, causing rolling around the longitudinal axis. (b) Rolling frequency measured at 100 fps for *N* = 216 sperm cells. (c) A representative trajectory of sperm wandering in TALP. (d) Circling motility in TALP + 1% PAM + 6.0 mM caffeine. These trajectories are recorded at 15 fps. Both real-time (transparent lines) and smoothed (solid lines) trajectories are shown. (e) MSDs for wandering and circling motility as a function of time. (f) The swimming behavior space for wandering (black) and circling (blue) phases. Purple dots (as control) represent data for circling sperm in TALP + 1% PAM without caffeine stimulation.

### Wandering motion in a Newtonian fluid

When stimulated with hyperactivation agonists, the flagellar beating pattern becomes highly asymmetric, leading to stochastic dynamics in TALP, resembling features of a persistent random walk (PRW)^53^. To quantify this wandering motion, we induced hyperactivation with 6.0 mM caffeine and extracted sperm trajectories over long durations (5–8 minutes, Fig. 1**c**, Movie S2). We then computed the mean-square displacement (MSD) ⟨|Δ***r***(*t*)|^2^⟩ from the trajectories with Δ***r***(*t*) = ***r***(*t*) − ***r***(0) being the displacement between the initial ***r***(0) and current position ***r***(*t*). A representative example is shown in Fig. 1**e**, exhibiting short-time ballistic behavior ∼ *t*^2^, followed by diffusive behavior 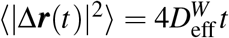, with an effective diffusivity of 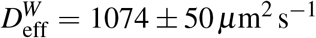 at long times. Note that at very long times, the MSD saturates, reflecting the finite size of the microfluidic chamber. The diffusive regime is induced by sudden reorientations at randomly distributed times. We found that on average, the sperm swam at a net translational speed of *U* = 20 ± 4 *µ*ms^−1^ and with a persistence time of *τ* = 3.9 ± 2.2 s (N = 20). Describing the motion as PRW in two dimensions, the diffusivity obeys 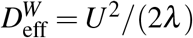, with *λ* = *D*_rot_ + *τ*^−1^. In agreement with previous reports^53^, this wandering motion was dose-dependent, with *U* = 80 ± 27 *µ*ms^−1^ and *τ* = 7.5 ± 5.8 s (N = 20) when stimulated with 3.0 mM caffeine. We further quantified wandering motion (6.0 mM caffeine) at the population level (N = 358) and found a large distribution of translation speeds ranging from *U* = *𝒪*(1) to *U* = *𝒪*(100) *µ*ms^−1^, which implies a large variation in 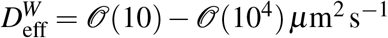(Fig. 1**f**). These results reflect a large behavioral variability across cells in a population coming from an individual sample.

### Circling motion in complex fluids

Upon entering a chamber filled with TALP + 1% PAM medium, sperm rolling becomes suppressed and the cells follow circular trajectories with large radii, which stem from slight asymmetries in the beating patterns^41, 53^. However, the addition of 6.0 mM caffeine to this mucus-mimicking complex fluid results in highly asymmetric flagellar beating patterns, and the sperm exhibit circling with much smaller radii (Fig. 1**e**, Movie S2). A typical MSD is shown in Fig. 1**e**, displaying three distinct features: persistent ∼ *t*^2^ behavior at short times, oscillatory dynamics at intermediate times—indicative of circling—and diffusive ∼ *t* scaling at long times, with an effective diffusivity of 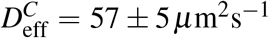. This circling motion has been studied theoretically, with predictions yielding an effective diffusivity^54, 55^ of 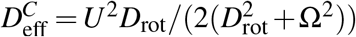, which decreases with increasing rotational speed (Ω). We quantified sperm circling at the population level with (N=418) and without (N=199) treatment with 6.0 mM caffeine (Fig. 1**f**), and measured swimming parameters including *U*, Ω, and 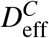. We notice that caffeine treatment increases Ω from 0.4 ± 0.3 s^−1^ to 2.6 ± 0.7 s^−1^, which led to a substantial decrease of 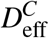 from 440 ± 147 *µ*m^2^s^−1^ to 174 ± 68 *µ*m^2^s^−1^. We also found that the population-level average of 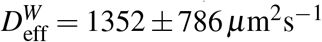 is an order of magnitude higher than that for circling. Therefore, the transition from wandering in a Newtonian fluid to circling in a complex fluid significantly limits spatial exploration while enhancing local exploitation.

### Circling-and-wandering in complex fluids

Under stimulation with 6.0 mM caffeine, the suppression of rolling in TALP + 1% PAM medium was not permanent for all sperm cells. We observed a subpopulation of cells that intermittently switched between non-rolling and rolling states, resulting in a mixed motility alternating between circling and wandering periods. To accurately characterize this mixed motility and obtain reliable statistics, we conducted new experiments in which sperm samples were further diluted 1:10 and injected into the microfluidic chip in a controlled manner. This approach allowed us to reduce the number of cells in the chamber, observe the dynamics of individual cells, minimize cell-cell interactions in the chamber, and reduce possible rheological heterogeneity that may arise at high cell densities. Under these conditions, we tracked 28 sperm after they entered the chamber. Rolling was suppressed in all of them upon entry, and they initially exhibited circular motion. However, 12 out of the 28 cells resumed rolling after a while, leading to a switch to wandering motion, followed by intermittent rolling suppression at later times, resulting in repeated transitions between circling and wandering periods (Fig. 2**a**, Fig. S2, Movie S3). Such transitions were evident in the rapid changes in Ω from zero to nonzero values and vice versa (Fig. 2**b**, Fig. S2). The average times spent in the circling and wandering periods were *τ*_*C*_ = 117.3 ± 88.8 s and *τ*_*W*_ = 84.8 ± 33.4 s (*N* = 12), respectively, and Ω remained constant across different constituent circular periods (Fig. 2**b**, Fig. S2). Consistent with the statistical picture obtained for pure circling and wandering, *D*_eff_ inferred for the constituent circling periods was significantly smaller than that of the constituent wandering periods (Fig. S3).

**Figure 2.**
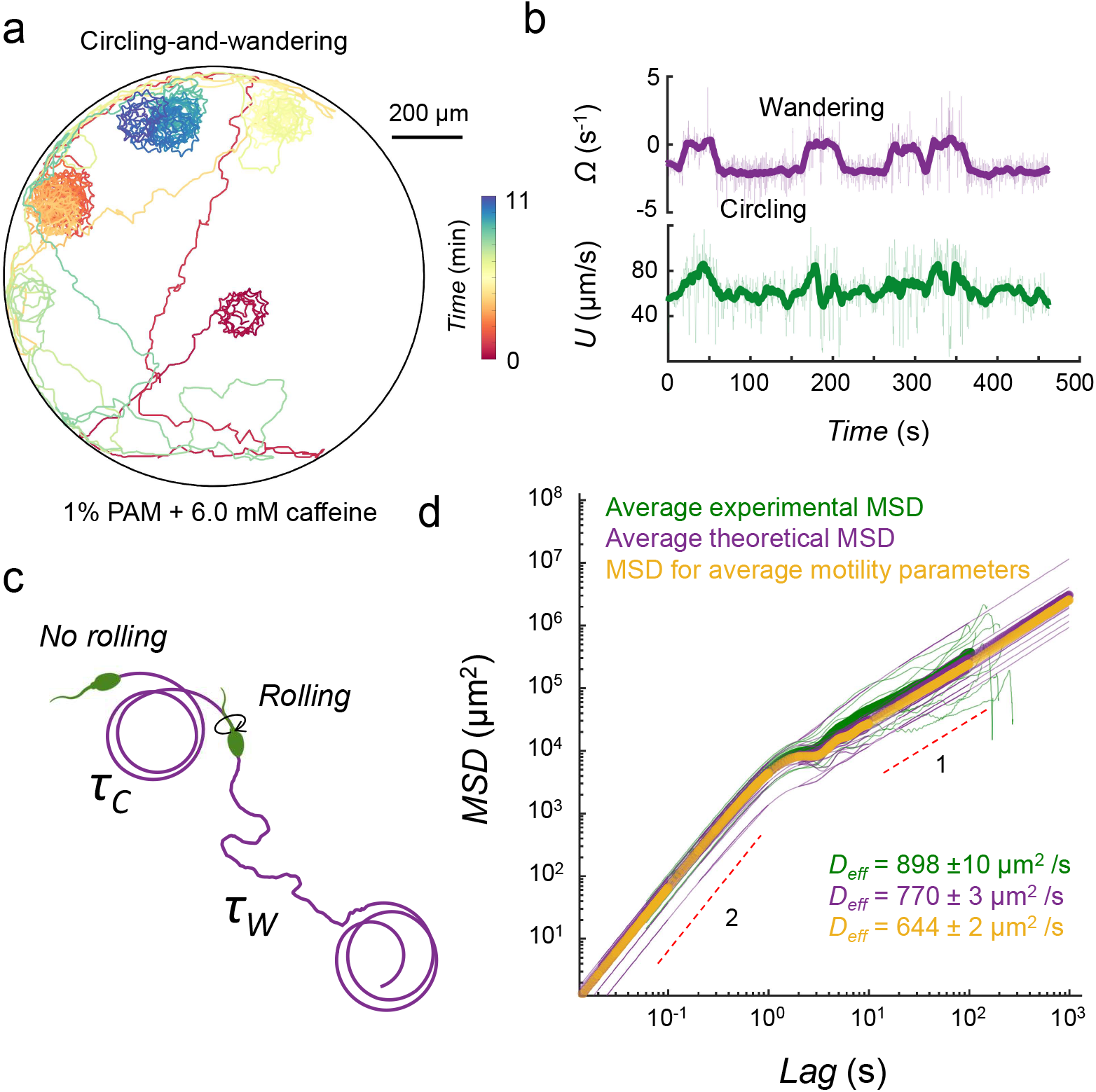
Sperm circling-and-wandering. (a) A representative trajectory of a circling-and-wandering sperm. (b) Instantaneous translational *U* and rotational Ω speeds corresponding to the trajectory in (a). (c) Sketch of the model system, where *τ*_*C*_ and *τ*_*W*_ denote the average time sperm spend in each phase, respectively. (d) Experimental measurements and theoretical predictions for the MSD. Motility parameters measured from the velocity correlation function of individual trajectories have been used as input for the theory in the purple curve. The yellow curve corresponds to the theoretical MSD obtained from the motility parameters averaged over all individual trajectories.

We also quantified sperm persistence length during wandering periods as the average distance over which the cell maintains its direction before changing it due to rotational fluctuations. We observed that this persistence length was significantly greater (*l*_p_ = 823 ± 587 *µ*m) compared to pure wandering in TALP (Newtonian fluid) under the same 6.0 mM caffeine treatment (*l*_p_ = 73 ± 40 *µ*m). This persistence was similar to that observed in pure wandering in TALP with milder caffeine stimulation (3.0 mM, *l*_p_ = 549 ± 405 *µ*m), but still significantly lower than the persistence length of progressive motion in TALP (*l*_p_ = 2110 ± 1268 *µ*m).

To investigate circling-and-wandering theoretically, we used a renewal theory^56, 57^ (see Methods), which provided a prediction for the MSD and the long-time effective diffusivity:

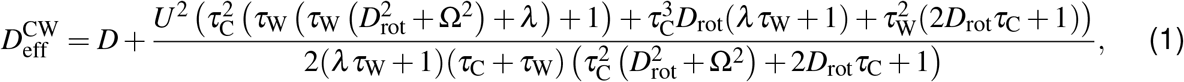

where *D* is the Brownian translational diffusivity and *λ* = *D*_rot_ + *τ*^−1^. This expression recovers coefficients for the pure individual phases (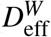 and 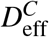) by setting *τ*_*C*_ = 0 (*τ*_*W*_ = 0) and taking the limit of *τ*_*W*_ → ∞ (*τ*_*C*_ → ∞), respectively (Fig. **S9b**). The MSD (with measured motility parameters as inputs) agrees with our experimentally measured MSD, averaged over all trajectories, thus confirming that the coarse-grained theory describes the observed trajectories and serves as a model for future investigations (Fig. 2**c** and **d**). A comparison of *D*_eff_ across all phases of motility shows that while pure wandering has higher effective diffusivities than pure circling, circling-and-wandering lies in between, suggesting its potential to bridge large-scale exploration and local exploitation strategies.

### Sperm-wall interactions: from concave to convex boundaries

To study sperm migration in complex geometries, we first quantified their swimming dynamics near concave boundaries (sidewall of the chamber) and convex (circular) pillars with different radii (*R* = 150 *µm* and *R* = 50 *µm*) under the environmental conditions discussed earlier.

#### Sliding of progressive sperm along boundaries

Near concave boundaries, progressive sperm aligned with the boundary upon collision and swam along it until the end of the measurement, consistent with previous observations^4^ (Fig. 3**a**). A similar pattern was observed near the circular pillars. In particular, sperm swimming directions were rectified upon collision with pillars; consequently, they swam along the pillar for only a short arclength, which was shorter for smaller pillars (N = 39, Fig. 3**b**). By measuring the cumulative changes in the direction of sperm swimming *θ*_cumulative_, i.e., the overall deflection angle Δ*θ* (Fig. S5**a**), we found that pillars rectified the swimming direction in a deterministic manner, as the deflection angle and contact arclength were strongly correlated (Fig. 3**c**).

**Figure 3.**
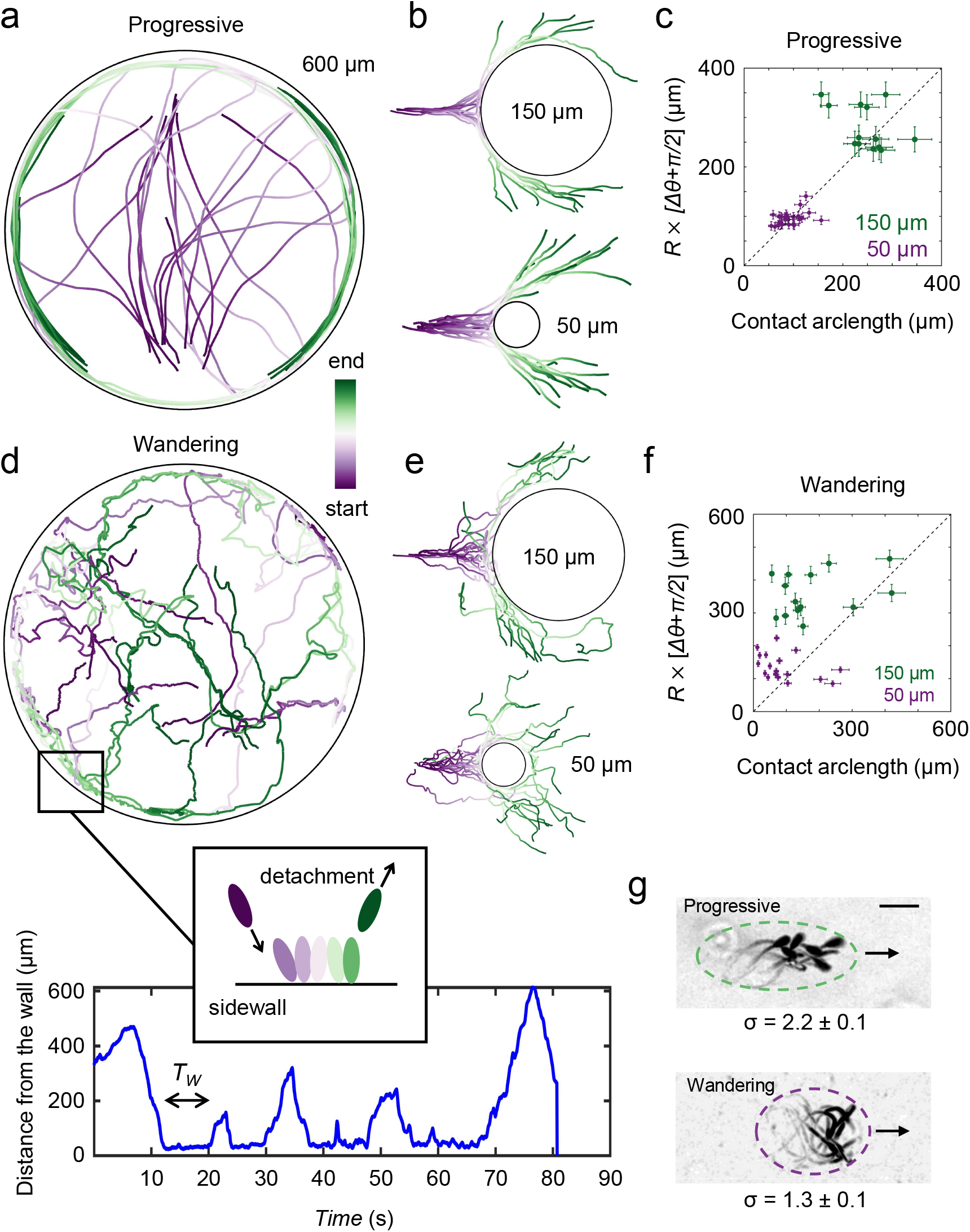
Sperm-wall interactions in the progressive and wandering phases. (a) Progressive motility in TALP allows for entrapment around the circular sidewall. (b) The swimming direction is deterministically rectified by pillars with *R* = 150 *µm* and *R* = 50 *µm*. (c) Sperm contact arclength on the pillar is linearly correlated with the deflection angle Δ*θ*. (d) The wandering phase allows sperm to explore the area enclosed by the circular sidewall. After each collision with the sidewall, sperm detach after a transitory dwell time *T*_*w*_. (e) Wandering sperm scatter from the pillars, with *R* = 150 *µm* and *R* = 50 *µm*, in a stochastic manner. (f) Sperm contact arclength on the pillar is not correlated with the deflection angle Δ*θ*. (g) Sperm aspect ratio in progressive and wandering phases.

#### Scattering of wandering sperm from boundaries

The interaction of wandering sperm (TALP with 6.0 mM caffeine) with concave boundaries was significantly different from that of their progressive counterparts. They did not align with the sidewall but remained near the contact point until they were scattered away (Movie S4), swam through the chamber interior, and collided with the boundary at another point (Fig. 3**d**). We quantified this observation by measuring the distance between the sperm and the sidewall, as well as the distribution of dwell times, which had an average of *τ*_dwell_ = 13.8 ± 14.8 s (*N* = 59, Fig. S5**e**).

Similarly, wandering sperm remained near the contact point when they collided with the convex pillars until they scattered away (Fig. 3**e**, Fig. S5**b**,**c**, and Movie S5). Unlike progressive sperm, the contact arclength on the pillar was not correlated with the deflection angle (Fig. 3**f**), and the relative standard deviation of the deflection angle was higher than that of progressive sperm (N = 73, Fig. S4**d**), highlighting the ‘noisy’ scattering of wandering sperm from the pillars.

In general, the random nature of wandering motility generates substantial orientational fluctuations that prevent sperm from aligning with boundaries. Furthermore, previous models suggested that the aspect ratio of a microswimmer determines its alignment dynamics when it collides with a boundary^58, 59^. We then measured the aspect ratio, *σ* = *W/L* (with width *W* and length *L*), of progressive (*σ* = 2.3 ± 0.2) and wandering (*σ* = 1.3 ± 0.2) sperm which revealed that the effective shape of the microswimmer changed from an ellipse in the progressive phase to a more spherical shape in the wandering phase (Fig. 3**g**, Fig. S5**a**,**b**). Therefore, wandering sperm are expected to experience less aligning torque from the sidewall (Fig. S5**c**). We note that the lower aspect ratio during wandering results from changes in the shorter wavelength and higher amplitude of the beating pattern.

#### Entrapment of circling sperm by convex boundaries

We then studied the interaction of the circling phase with pillars in TALP + 1% PAM. Without caffeine treatment, sperm swam in large circles and interacted with *R* = 150 *µm* pillars. Typically, when the curvature of the swimming path was positive (with respect to the pillar) at the contact point, the sperm became trapped; otherwise, they scattered from the pillar. Local contacts led to the alignment of the swimming direction along the pillar’s periphery, resulting in persistent entrapment. The first entrapment event occurred within 0.5–1 minute of sperm entry into the chamber, with up to ∼ 30 and ∼ 100 cells becoming trapped within 3 and 10 minutes, respectively (Fig. 4**a**, Movie S6). The trapped sperm exhibited nonuniform chirality, with 85 ± 11% displaying counterclockwise (CCW) motion (Fig. S6**a**). We also noticed that temporary rolling events at the contact point reversed the swimming chirality, and consequently transitioned sperm-wall interactions from scattering to entrapment, highlighting that having positive curvature at the contact point is key for entrapment (Fig. 4**b**, Movie S7).

**Figure 4.**
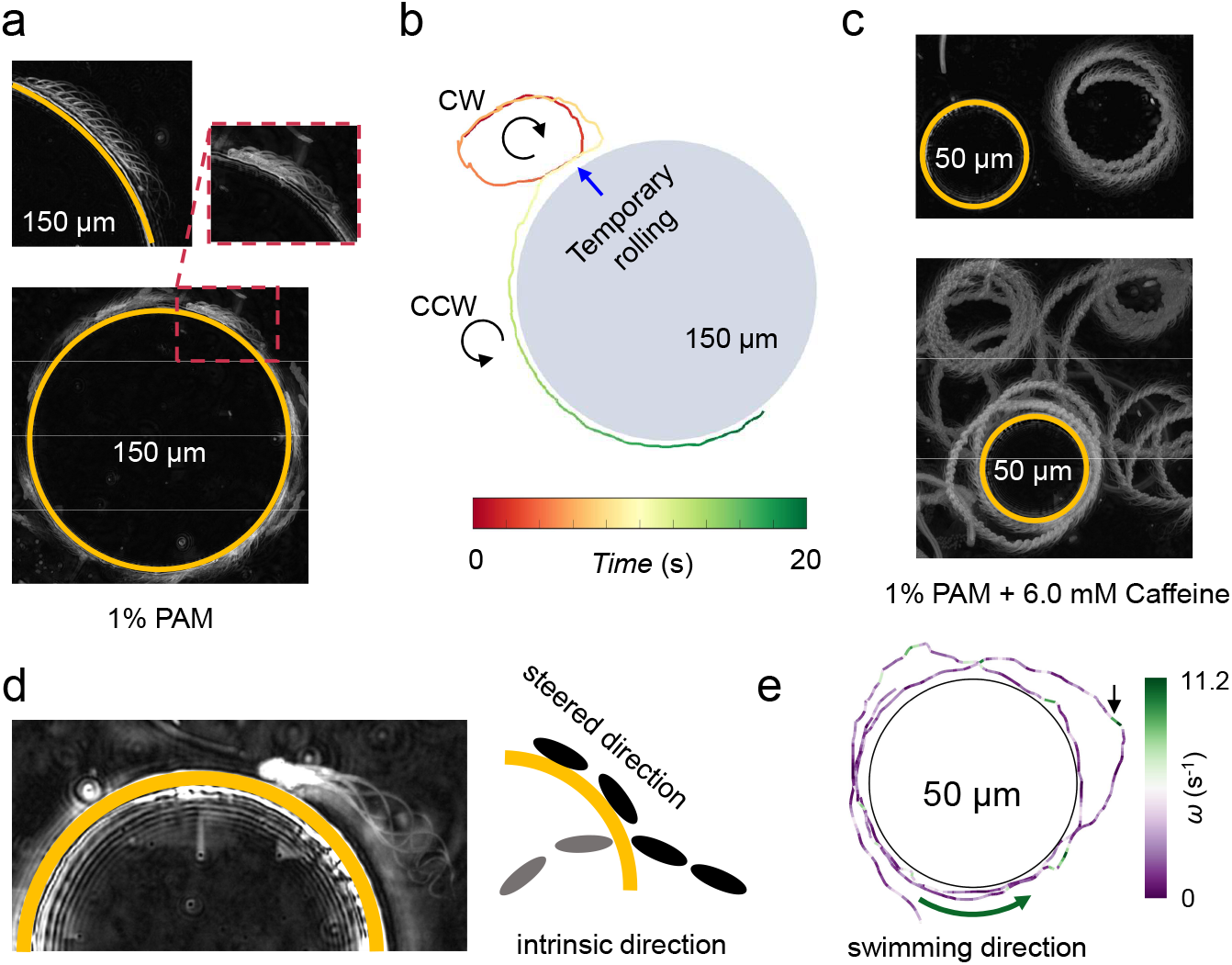
Sperm-wall interactions in the circling phase. (a) Without hyperactivation, sperm in the circling phase feature large radii and become trapped around pillars with *R* = 150 *µm*. Entrapment of the first sperm (top) occurs within 1 minute of entry into the chamber, increasing to 30 sperm within 3 minutes. (b) Temporary rolling events can change the chirality of circling motility, allowing transitions from scattering to entrapment or vice versa. (c) With hyperactivation, circling motility features tight circles, and sperm become trapped around *R* = 50 *µm* pillars. (d) When trapped, the sperm’s intrinsic swimming direction is not necessarily aligned with the steered swimming direction, reducing its rotational speed. (e) A representative trajectory of a sperm circling around a pillar, color-coded with respect to its rotational speed. The arrow indicates a high rotational speed (deep bending in the flagellum) when the sperm is not in contact with the pillar.

However, a positive curvature at the contact point alone was insufficient for entrapment. Sperm with high-radius circular paths scattered from *R* = 150 *µ*m pillars after partial rectification (Fig. S7**a**). Our measurements suggest that the maximum trajectory radius for entrapment was 520 ± 57 *µ*m, and the minimum was 92 ± 11 *µ*m (Fig. S7**b**). Without caffeine treatment, the circling sperm did not show a significant interaction with the *R* = 50 *µ*m pillars within the 10-minute observation period, indicating that the ratio between the radius of the pillar and the radius of circling is crucial for entrapment. To quantify these observations, we measured the radii of circular trajectories without caffeine (Fig. S7**c**) and found that the average radius (1.2 ± 0.7 mm) exceeded significantly *R* = 50 *µ*m, explaining the lack of interaction with smaller pillars. However, a subset of cells had trajectories within the trapping range (*>* 92 *µ*m and *<* 520 *µ*m) for *R* = 150 *µ*m pillars, accounting for approximately 11.3% of the cells, the estimated fraction of trapped sperm.

With 6.0 mM caffeine treatment, circling sperm did not become trapped around *R* = 150 *µm* pillars. This is not surprising, as after measuring the radii of the circling trajectories at the population level (Fig. S7**d**), we found that the average radius (36 ± 10 *µm*) was below the minimum entrapment radius (92 ± 11 *µm*) for the pillars *R* = 150 *µm*. In contrast, and as expected, entrapment occurred around the *R* = 50 *µm* pillars in 10 minutes (1–3 cells per pillar, Fig. 4**c**, top at *t* = 0 and bottom at *t* ≈ 10 min, Movie S6). To further quantify this observation, we measured the velocity correlation function for *N* = 7 sperm before and after entrapment to determine their rotational speed. We found that Ω decreased significantly upon entrapment (Ω_trapped_ = 1.4 ± 0.1 s^−1^ vs. Ω_free_ = 2.0 ± 0.5 s^−1^, *p*-value = 0.01; Fig. S8), likely due to misalignment between the intrinsic swimming direction and the boundary constraint (Fig. 4**d, e**).

### Sperm migration in a complex environment

Having established an experimental classification of hyperactivated motility in Newtonian and complex fluids, we investigated sperm transport in complex media using stochastic simulations.

Different phases of sperm motility (Fig. 5**a**) are described according to Sec. 2.1, using experimentally measured parameters as input (see Methods, Table 1). The complex environment is modeled as a two-dimensional porous medium composed of overlapping discs of radius *R*_0_, reflecting the quasi-2D, crowded nature of the folds of the FRT. We keep a constant packing fraction 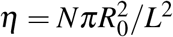 (with number of obstacles *N* and system size *L*) and vary the disc radius *R*_*0*_^60^. This generates porous media with varying average chord lengths, *l*_*c*_ = *πR*_0_*/*(2*η*), and thus different numbers of corners and dead-end passages (Fig. 5**c**).

**Figure 5.**
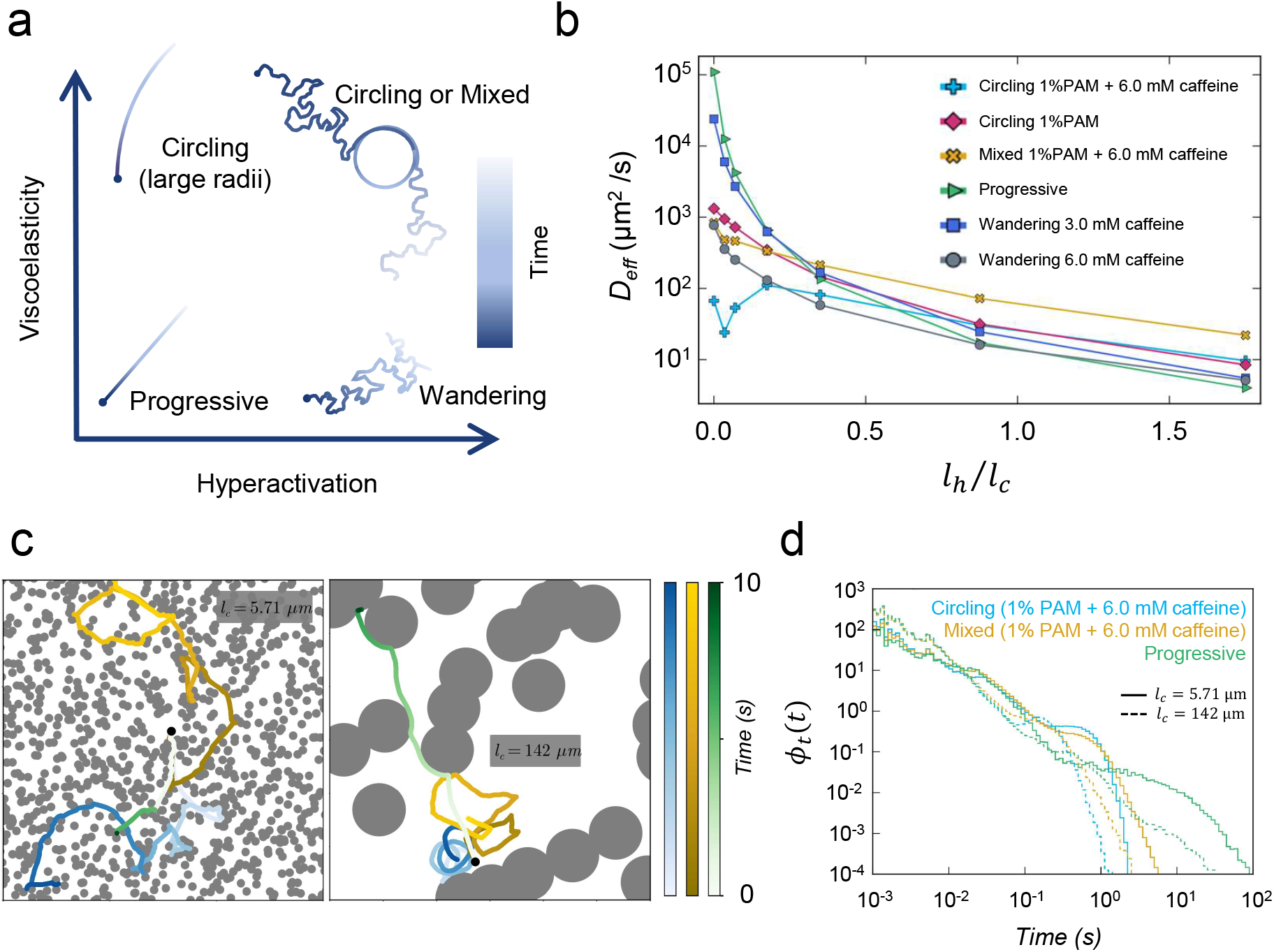
Sperm motion in porous media. (a) Different swimming behaviors promoted by hyperactivation and the viscoelasticity of the swimming medium. (b) Effective diffusion coefficient (*D*_eff_) measured for sperm moving in porous media under different conditions. Here, *l*_*h*_ denotes the sperm head length and *l*_*c*_ represents the average chord length. (c) Typical trajectories of progressive (green), circling (blue), and mixed-phase (yellow) motion in two different porous media. The obstacle radii are 2*µ*m and 50*µ*m in the left and right panels, respectively. Color coding corresponds to elapsed time. (d) Trapping time distributions, *φ*_*t*_(*t*), for the three phases in the same environments as in (c).

**Table 1.** Experimentally-measured averaged motility parameters of sperm in different phases. These are used as input for our simulations. ‘n.a.’ denotes ‘not applicable’.

| motility pattern (fluid, chemical) | $U$ ($\mu\text{ms}^{-1}$) | $D_{\text{rot}}$ ($\text{s}^{-1}$) | $\tau$ (s) | $\Omega$ ($\text{s}^{-1}$) | $\tau_C$ (s) | $\tau_W$ (s) |
| --- | --- | --- | --- | --- | --- | --- |
| progressive (Newtonian, not stimulated) | 101.3 | 0.047 | n.a. | n.a. | n.a. | n.a. |
| PRW (Newtonian, 3 mM caffeine) | 79.8 | 0.047 | 11.63 | n.a. | n.a. | n.a. |
| PRW (Newtonian, 6 mM caffeine) | 19.9 | 0.047 | 4.78 | n.a. | n.a. | n.a. |
| chiral (1% PAM, not stimulated) | 58.22 | 0.14 | n.a. | 0.4 | n.a. | n.a. |
| chiral (1% PAM, 6 mM caffeine) | 71.24 | 0.14 | n.a. | 2.3 | n.a. | n.a. |
| circling/wandering (1% PAM, 6 mM caffeine) | 82.11 | 0.19 | 0.67 | 1.906 | 66.67 | 31.25 |

We quantified sperm dynamics by measuring the MSD, which grows linearly with time at long times, ⟨|Δ***r***(*t*)|^2^⟩ = 4*D*_eff_*t*, allowing us to extract *D*_eff_ as a measure of the dispersion of cells in the porous medium (Fig. S9**a** and **c**). In dilute media (corresponding to large average chord lengths *l*_*c*_), our simulations show that *D*_eff_ is largest for progressive and smallest for circling sperm (1% PAM + 6.0 mM caffeine), while the mixed phase lies in between, confirming our previous statistical characterization of motility in free space (Fig. 5**b**). Decreasing the average chord length *l*_*c*_ leads to a stark decrease in *D*_eff_ for the progressive phase by up to four orders of magnitude, making the progressive phase the least effective in a porous medium. The effective diffusivity *D*_eff_ also decreases for the other phases, but the effect is not as pronounced. Interestingly, our results further show that the mixed phase is only weakly affected by changes of the environment and becomes the most efficient transport strategy as the average chord length is decreased.

This behavior can be rationalized by inspecting the trajectories in a medium with a long average chord length, *l*_*c*_ = 142 *µ*m (Fig. 5**c**). Here, progressive sperm are able to explore large areas through their almost straight swimming motion, while circling sperm circle in the open pore space without migrating far from the initial position. During the circling-and-wandering phase, sperm can explore the environment more than those with pure circling, but their continuous random orientational changes lead to an overall smaller displacement than that of the progressive phase. This behavior changes drastically when the average chord length is decreased to *l*_*c*_ = 5.71*µ*m (which is comparable to the sperm head size), where progressive sperm remain stuck in the corners of the porous matrix, while the other motility patterns lead to larger exploration of the pore space. We emphasize that the mixed phase remains more efficient than the pure circling, as the wandering periods helps the cell move farther.

These observations can also be quantified by measuring the trapping times, i.e., the time sperm are trapped at obstacle boundaries and in narrow channels where they barely move (Fig. 5**d**). The distributions show distinct features for the different phases and indicate long trapping phases for progressive sperm. For the mixed and circling phases, the trapping times are shorter and the decay of the distribution is faster. This is due to cells being able to reorient, which allows them to escape from tight corners and boundaries, thus reducing the trapping times^61, 62^. From our simulations results, we may conclude that hyperactivated motility, regardless of fluid properties, shows enhanced dispersion copmpared to progressive motility; that is, randomness in swimming behavior is crucial for migration in confined and complex geometries, whereas progressive motility is more suitable for functional regions with less complexity.

In summary, our results show that hyperactivation enables sperm to dynamically adjust their motility, producing distinct motion phases depending on the viscoelasticity of the fluid. At low viscoelasticity, hyperactivated sperm display wandering behavior, while at high viscoelasticity, they shift to circling or circling-and-wandering modes. Wandering sperm, less affected by geometric constraints, promote random exploration in complex FRT environments. In contrast, circling sperm, characterized by lower effective diffusivity, favor localized exploitation, often being trapped around pillars. This may allow sperm to exploit viscoelastic niches and remain near targets such as the oocyte. The circling-and-wandering motility, therefore, represents an adaptive swimming strategy that allows for a trade-off between exploring large areas and exploiting localized regions (Fig. 6), potentially enhancing search efficiency, similar to other intermittent search strategies observed across various biological systems^63, 64^. Our simulations further indicate that, as geometrical complexity increases, circling-and-wandering exhibits higher effective diffusivity than wandering or circling alone, suggesting improved dispersion due to reduced trapping and enhanced movement through pores during the circling and wandering periods, respectively.

**Figure 6.**
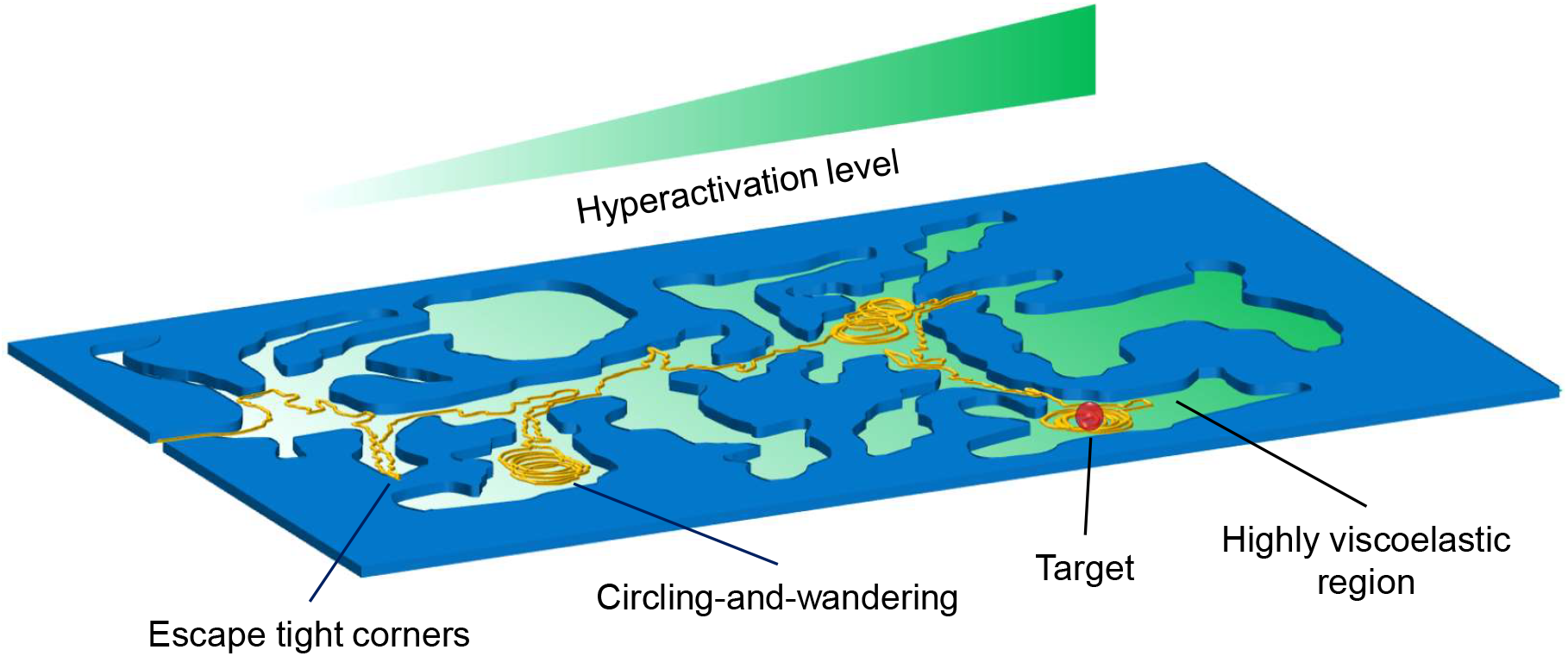
Schematic of sperm circling-and-wandering migration strategy within the FRT

One might argue that strong rheological heterogeneity in the medium could be sufficient to trigger transitions between wandering and circling motions. However, we emphasize that the observed circling-and-wandering occurred despite the medium’s near-homogeneous rheology, indicating that stochastic transitions between these modes are intrinsic to hyperactivation. Since drastic rheological changes in the upper FRT appear unlikely, circling-and-wandering could represent a key migration strategy in these regions and may provide a foundation for a model describing mammalian sperm chemotaxis, analogous to the well-established bacterial run-and-tumble model^56, 65^. We should stress that despite similarities between bacterial tumbling and sperm circling, a key difference exists: whereas bacterial tumbling occurs on the order of milliseconds (or a fraction of the run time), sperm circling unfolds over minutes, with a duration comparable to that of the wandering phase. Therefore, and as we suggested earlier, the circling phase functions as a local exploitation strategy rather than merely serving to change the swimming direction.

In the context of chemotaxis, a key question is how circling-and-wandering motion can be modulated—specifically, by controlling the transition frequency between constituent phases. To address this, it is crucial to investigate how sperm rolling is reversibly and stochastically suppressed in viscoelastic media. Based on previous studies of bacterial swimming in complex fluids^66^, we predict that fluid viscoelasticity reduces the angle between the sperm head and the flagellar beating plane, potentially by altering the conformation of the dynamic basal complex^67, 68^, which leads to the suppression of rolling. However, alternative mechanisms, such as increased viscous resistance or enhanced elastic stresses, may also contribute.

Although we used a chemical agonist to induce hyperactivation, recent evidence suggests that activation of CatSper and the resulting hyperactivation, can also be triggered by thermal cues^69^. This implies that hyperactivation is a shared behavior in both chemotactic and thermotactic responses. Therefore, our findings motivate future investigations into the role of circling-and-wandering in sperm thermotaxis, pointing to a potential unifying mechanism governing chemotactic and thermotactic navigation. Another important avenue for future research is understanding how directional cues such as chemical gradients, directional mucus flow within the oviduct, or gradients in the viscoelasticity of the fluid, potentially modulate circling-and-wandering and steer movement in a controlled manner. This study also motivates further theoretical investigation into how detailed hydrodynamic interactions and sperm morphology influence migration in geometrically complex environments. Our findings offer insights for the development of assisted reproductive technologies, particularly in selecting for hyperactivated motility to improve the efficacy of current techniques^70–72^.

## 1 Methods

### 1. Microfluidic Device

Our microfluidic device comprised a straight microchannel (width ∼ 400 *µ*m) connected to circular Hele-Shaw chambers on either side, each with a diameter of 1200 *µ*m. For wall-interaction experiments, we fabricated concentric pillars with diameters of 100 and 300 *µ*m within the chambers. The height of the device was 30 *µ*m throughout. The chambers were initially filled with the swimming media, e.g., TALP + caffeine or TALP + 1% PAM + caffeine. Later, the sperm sample was injected into the main channel using gravity, and by reducing the flow, sperm cells were able to swim inside the chambers without perturbations caused by external fluid flow. Microfabrication was carried out using standard soft lithography techniques.

### 1.2 Sperm Preparation and Chemicals

Commercially available cryopreserved bull sperm samples (5.5 to 6.5 years of age) diluted in egg yolk extender were generously donated by URUS Holding Company, Ithaca, NY. The ejaculate concentration was 50 × 10^6^ sperm per straw (0.25 mL) with a pre-freeze motility of 65%. A combination of gentamicin, tylosin, lincomycin, and spectinomycin was added to the semen as antibiotics prior to cryopreservation. For each experiment, two straws were thawed in a 38^°^C water bath for 30 s, diluted with 1 mL TALP^**?**^, incubated at 38^°^C and 5% CO_2_ (Thermo Fisher) for 30 minutes, and then used immediately. The sperm samples and microfluidic device were maintained at 38^°^C throughout the experiments using a heated glass microscope plate (Bioscience Tools). The following recipe was used for making TALP: NaCl (110 mM), KCl (2.68 mM), NaH_2_PO_4_ (0.36 mM), NaHCO_3_ (25 mM), MgCl_2_ (0.49 mM), CaCl_2_ (2.4 mM), Hepes buffer (25 mM), glucose (5.56 mM), pyruvic acid (1.0 mM), penicillin G (0.006% or 3 mg/500 mL), and bovine serum albumin (20 mg/mL). Long-chain PAM with a molecular weight of 5–6 MDa was added to make the standard medium viscoelastic.

### 1.3 Microscopy

The sperm hyperactivated motility inside the circular chambers was observed and recorded via a Zeiss phase-contrast microscope (10 × objective) equipped with a Neo sCMOS high-speed digital camera (Andor). NIS Elements imaging software (Version 4.0; Nikon) was used for image acquisition. The rolling of bull sperm can be observed under phase optics, as the light intensity of the paddle-shaped head varies with rolling. To minimize the effect of ambient temperature changes on sperm motion, the microfluidic devices were kept at 38^°^C using a heated glass slide (Bioscience Tools).

### 1.4 Image Processing

The acquired videos were processed to extract sperm trajectories using TrackMate and a custom MATLAB code. Each trajectory was coarse-grained by averaging every five consecutive points to remove rapid oscillatory motion in the sperm head. Using the smoothed trajectories, we then calculated the translational velocity, ***V*** (*t*), and the normalized velocity autocorrelation function:

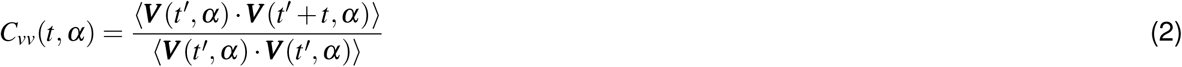

where *α* = 5 was used as a coarse-graining parameter. This velocity correlation function was then used to estimate values for rotational speed and relaxation times. We also measured the sperm reorientation angle,

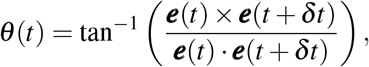

with the cumulative angle given by 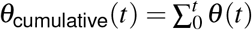.

After demonstrating that wandering, circling, and mixed phases are all diffusive in nature, we inferred diffusion coefficients based on the MSD, or on rotational and translational speeds and relaxation times for trajectories shorter than 150 points.

### 1.5 Theory and Simulation

#### 1.5.1 Theoretical derivation of the effective diffusivities for the mixed phase: A renewal approach

Here, we first discuss the MSDs of the two different phases: ⟨|Δ***r***(*t*)|^2^⟩ _*C,W*_ = ⟨|***r***(*t*) − ***r***(0)|^2^⟩ with initial position ***r***(0) and noise average ⟨·⟩. We note that the MSDs for the circling^54, 55^ and wandering phases^73^ have been previously reported and we refer the reader to previous work for a derivation. The MSD for the circling phase, where the sperm is modeled as an active Brownian circle simmer, evaluates to^54, 55^

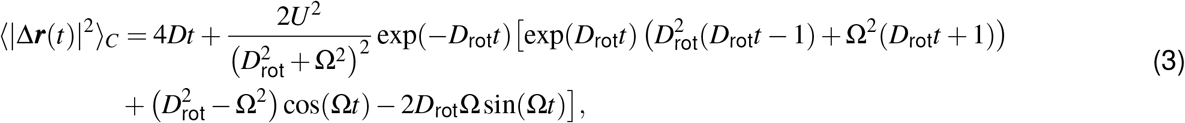

with swim speed *U*, angular velocity Ω, and rotational and translational diffusivities, *D*_rot_ and *D*, respectively. The wandering phase is modeled as a persistent random walk, where a particle randomly changes its swimming direction at exponentially distributed times, exp(− *t/τ*)*/τ* with average time between two events *τ*^73^. Thus, the MSD reads

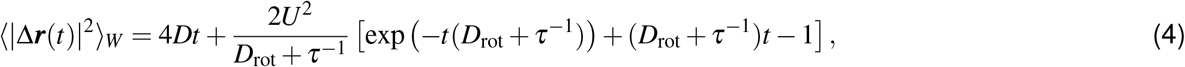

where the effects of translational and rotational diffusion are also included. We note that the contributions of rotational diffusion and the random change of swimming direction are additive at the level of the MSD and cannot be distinguished from one another. These two MSDs serve as input to compute the MSD in the mixed phase. The latter has not been derived before and, therefore, we present its derivation using a renewal framework.

The mixed phase consists of alternations of circling (*C*) and wandering (*W*) phases. We assume the agent spends an average time *τ*_C_ and *τ*_W_ in each of both phases, respectively, and that at the start of a new phase it moves along a new, random direction. A theoretical framework, allowing to couple two distinct phases, has been previously reported in terms of renewal equations^74^ and among others applied to study the run-and-tumble motion of bacteria in free space^56, 57, 75^ and their hop-and-trap dynamics in porous media^62^. Here, we denote by *P*(***r***,*t*) the probability density to be in a mixed phase at position ***r*** at time *t*. It can be decomposed as a sum of the probability to be in a circling phase *P*_*C*_(***r***,*t*) and a wandering phase *P*_*W*_ (***r***,*t*): *P*(***r***,*t*) = *P*_*C*_(***r***,*t*) + *P*_*W*_ (***r***,*t*). To characterize the circling phase we require two ingredients: the probability to move a distance ***r*** during time *t* in the chiral mode, 𝕡_*C*_(***r***,*t*), and the probability that the circling phase ends at time *t, ϕ*_*C*_(*t*) = exp(−*t/τ*_*C*_)*/τ*_*C*_. Similarly, for the wandering phase we introduce: the probability to move a distance ***r*** during time *t* in the persistent-random-walk mode, 𝕡_*W*_ (***r***,*t*), and the probability that the wandering phase ends at time *t, ϕ*_*W*_ (*t*) = exp(− *t/τ*_*W*_)*/τ*_*W*_. Assuming that the agents are in a non-equilibrium stationary state at time *t*, we need to account for the probability that the cells have spent their entire time up to *t* in *C* (*W*), respectively:

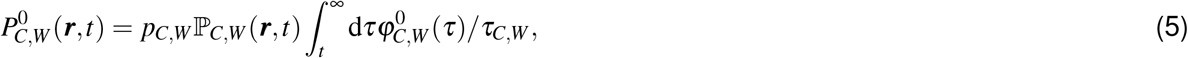

where *p*_*C*_ = *τ*_*C*_*/*(*τ*_*C*_ + *τ*_*W*_) and *p*_*W*_ = 1 − *p*_*C*_ denote the fraction of time spent in *C* and *W*, respectively. Further, 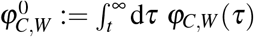 denotes the survival probability, i.e., the probability that the agent remains in *C* (*W*). The time integral in Eq. (5) corresponds to the probability density to switch from one to the other phase for the first time. See Ref.^57^ for more detailed explanations.

The two phases, *P*_*C*_(***r***,*t*) and *P*_*W*_ (***r***,*t*), are then described by the integral equations:

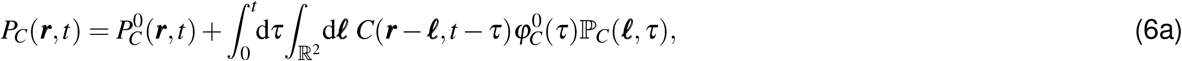

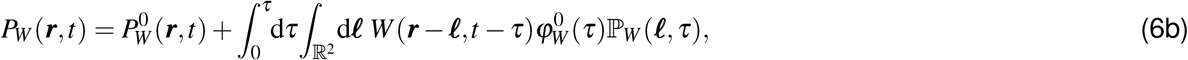

where we denote by *C*(***r***,*t*) and *W* (***r***,*t*) the probabilities (per unit time) to start a circling or wandering phase, respectively. Thus, the probability to be in C (W) at time *t* and position ***r*** is the sum of the probability to be in C (W) without ever having been in W (C) before and the sum over all previous changes between different phases, encoded in the integral. We note that neglecting 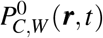 corresponds to a model where the cells start the C (W) phase at time *t*, which is rather hard to observe experimentally.

Next, we introduce equations of motion for the unknowns *C*(***r***,*t*) and *W* (***r***,*t*), which couple the two processes:

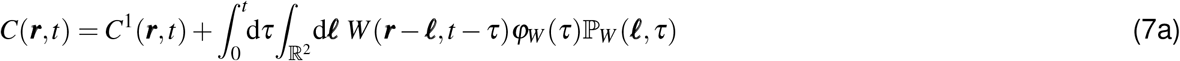

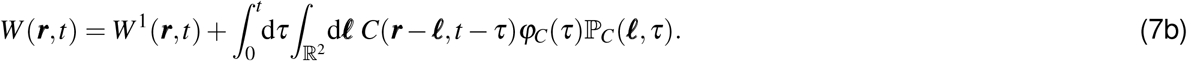

Here, we have again accounted for the probability to start a C (W) phase for the first time separately using

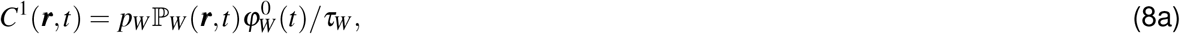

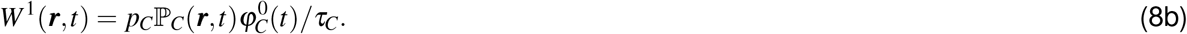

Moving to Fourier-Laplace space (***r*** → ***k*** and *t* → *s*) and using the convolution theorem, the set of equations has a formal solution^75^:

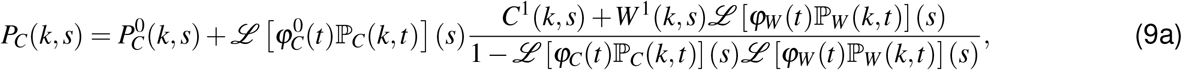

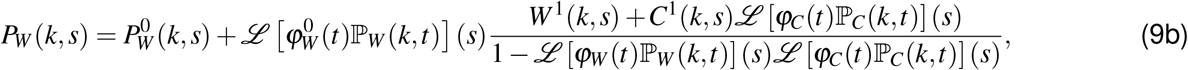

where 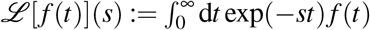 denotes the Laplace transform of a function *f* (*t*). Since the problem is translationally invariant, the quantities depend on the wavenumber *k* = |***k***| only. To obtain the MSD for the mixed phase, we are interested in the small-wavenumber expansion of the probability density, which, in Laplace space, assumes the form:

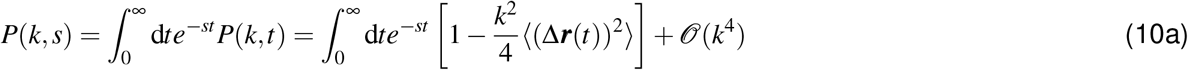

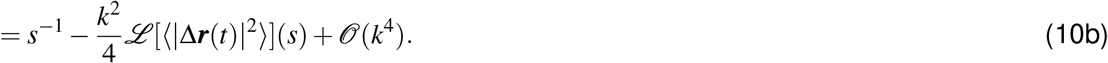

Therefore, we can also expand the propagators up to *𝒪*(*k*^4^) via

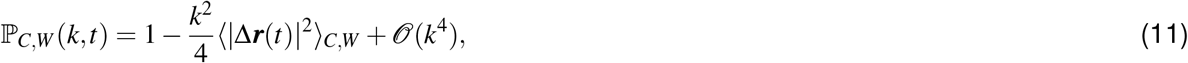

Inserting these expansions into Eqs. (9a)-(9b) and performing the Laplace transforms, allows calculating an analytical expression for *P*(*k, s*) to second order in *k* and hence the MSD in Laplace space, which can be transformed back to time analytically using Mathematica:

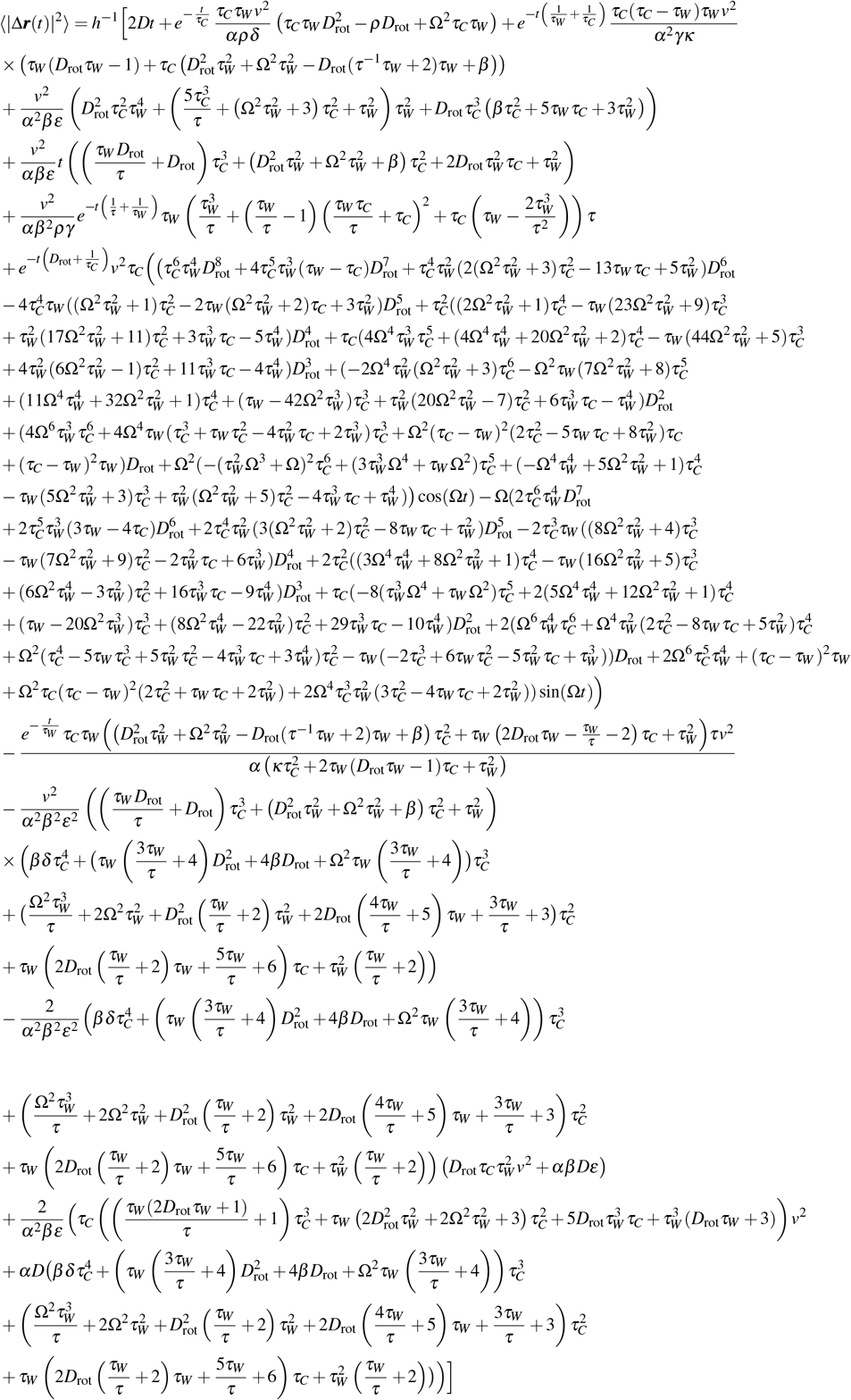

with

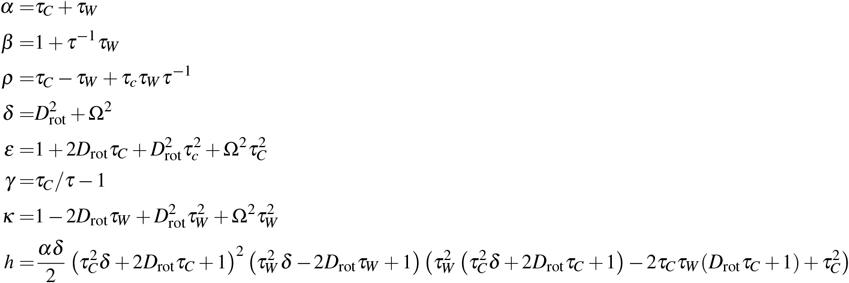

We obtain the long-time effective diffusivity for the mixed phase *D*^M^_eff_ by taking the limit

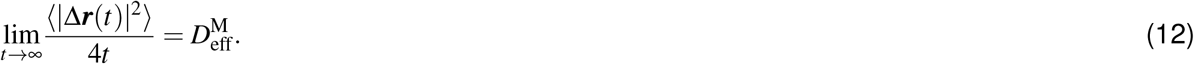

The latter is reported in the main text [Eq. (1)].

#### 1.5.2 Brownian dynamics simulations of active agents in porous media

To study transport of sperm cells in complex geometries, we simulate active agents, following the dynamics described above, in a two-dimensional Lorentz gas, where the obstacles are modeled as overlapping discs of radius *R*_0_ in a square domain of length *L*^76^. We ignore hydrodynamic interactions between the swimmers and the obstacles and perform Brownian dynamics simulations. The particles interact with the obstacles via a Weeks-Chandler-Anderson (WCA) potential:

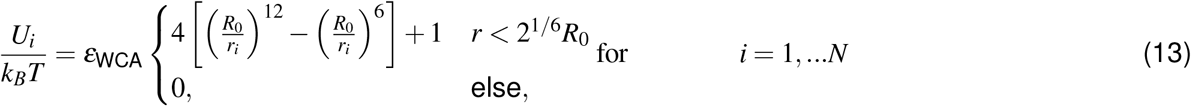

with the number of obstacles *N*, non-dimensional potential strength *ε*_*WCA*_, Boltzmann constant *k*_*B*_, and temperature *T*. Further, the distance between the particle and the *i*−th obstacle is *r*_*i*_ = |***r***(*t*) − ***r***_*i*_|, where ***r***_*i*_ denotes the position of the *i*−th obstacle. The equations of motion are:

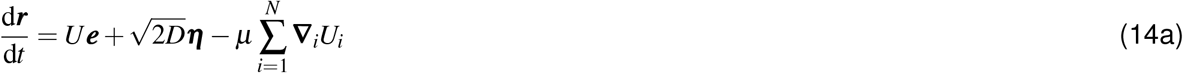

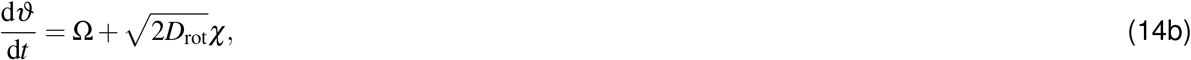

with mobility *µ* = *D/*(*k*_*B*_*T*). Further, ***e*** = (cos *ϑ*, sin *ϑ*) is the swimming direction and ***η***(*t*) and *χ*(*t*) are two independent Gaussian white noise processes of zero mean and delta-correlated variance, ⟨*η*_*i*_(*t*)*η*_*j*_(*t*^′^)⟩ = *δ*_*i j*_*δ* (*t* − *t*^′^) for *i, j* = 1, 2 and ⟨*χ*(*t*)*χ*(*t*^′^)⟩ = *δ* (*t* − *t*^′^), respectively. We simulate the wandering phase by setting Ω = 0 and changing orientation *ϑ* at exponentially distributed times ∼ exp(− *t/τ*)*/τ*. Finally, the mixed phase is simulated by changing from one phase to another at times ∼ exp(− *t/τ*_*C,W*_)*/τ*_*C,W*_ ; at the switching events, the agent starts moving along a new, random direction. We set the potential strength to *ε*_WCA_ = 100. We further choose parameter values for the active particles, which are extracted from the experimental observations (Table 1) and set the translational diffusivity to *D* = 0.01*µ*m^2^s^−1^.

To model the complexity of the medium, we vary the radius *R*_0_ ∈ [1, 50]*µ*m of the obstacles by keeping the packing fraction of the obstacles fixed, 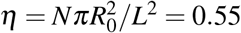 (with *N* = 7000). The latter leads to porous media that have varying mean chord length (*l*_*c*_ = *πR*_0_*/*(2*η*)), ranging from *l*_*c*_ ∈ [2.86, 143]*µ*m.

For the simulation, we discretize the equations of motion for the chiral phase according to the Euler-Maruyama scheme:

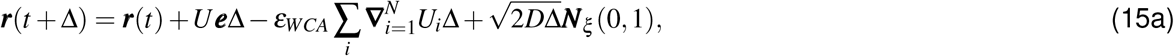

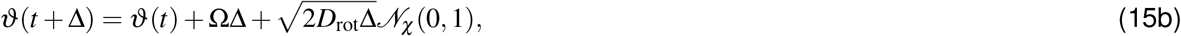

where ***N***_***ξ***_ (0, 1) and *𝒩*_*χ*_ (0, 1) are independent, normally distributed random variables of zero mean and unit variance. The time step is chosen as Δ = 10^−4^ and 10^3^ trajectories are obtained up to a total time of 10^4^. The simulation was programmed in Julia using neighbor lists for efficiency^77^.

To compute the trapping time distributions *φ*_*t*_(*t*) we measure the instantaneous displacement 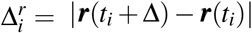 for 80 trajectories and apply a median filter to smoothen the time-series data. Trapping phases are then classified using the criterion that 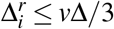. The long tails of the trapping-time distributions do not change significantly by varying the threshold. We consider a trapping event if its length exceeds 10Δ.

## 2 Acknowledgments

M.Z. thanks Ned S. Wingreen for insightful discussions about sperm circling-and-wandering and Susan S. Suarez for stimulating conversations about sperm hyperactivation that led to this project. C.K. acknowledges helpful discussions with S. Mandal and P. Das.

## 3 Supplementary figures

**Figure S 1.**
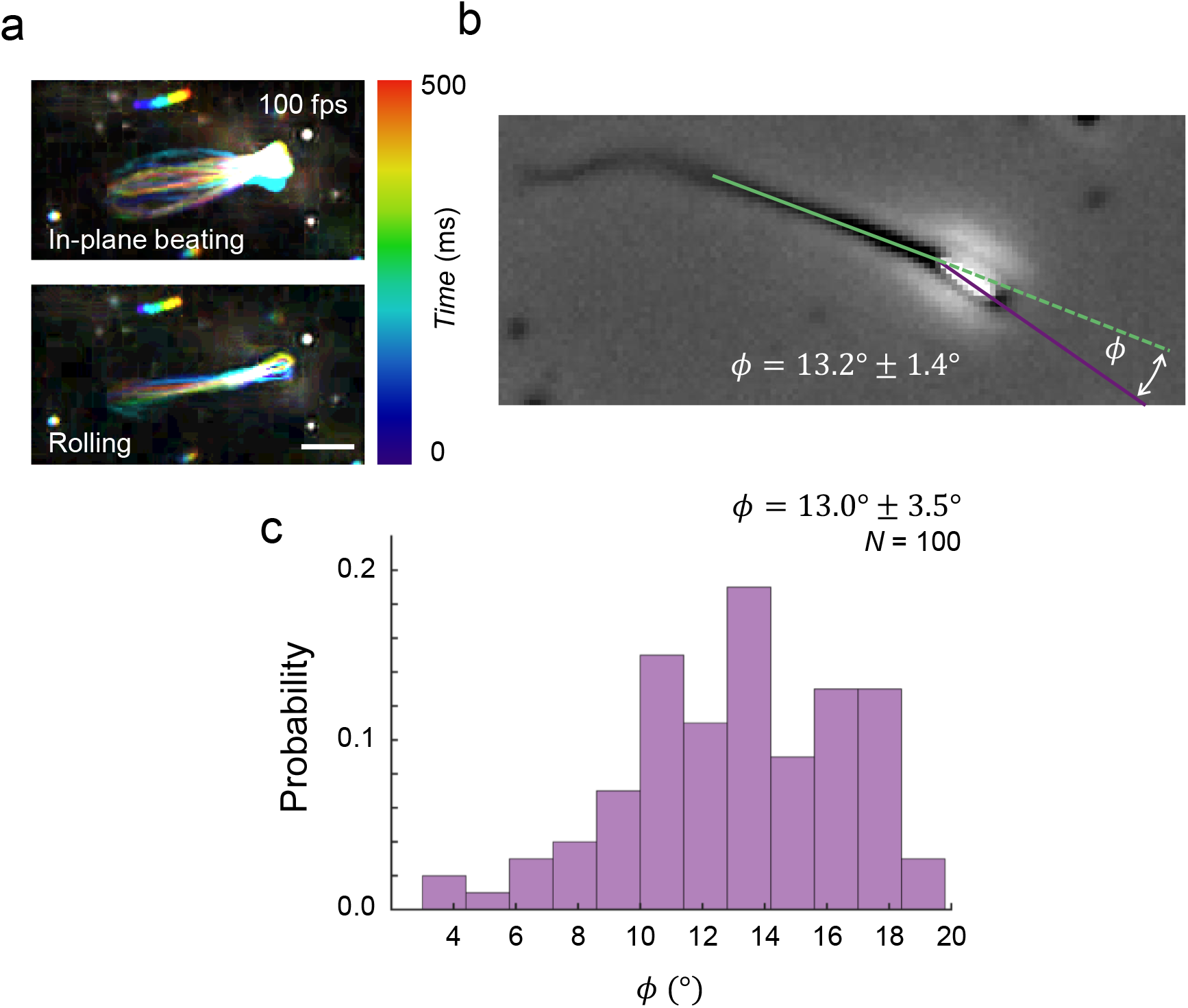
Sperm head morphology and rolling. (a) Bull sperm motility observed at 100 fps under phase-contrast microscopy. (b) Bull sperm head tilted out of the flagellar beating plane. (c) Histogram of the head angle with respect to the beating plane for *N* = 100 sperm.

**Figure S 2.**
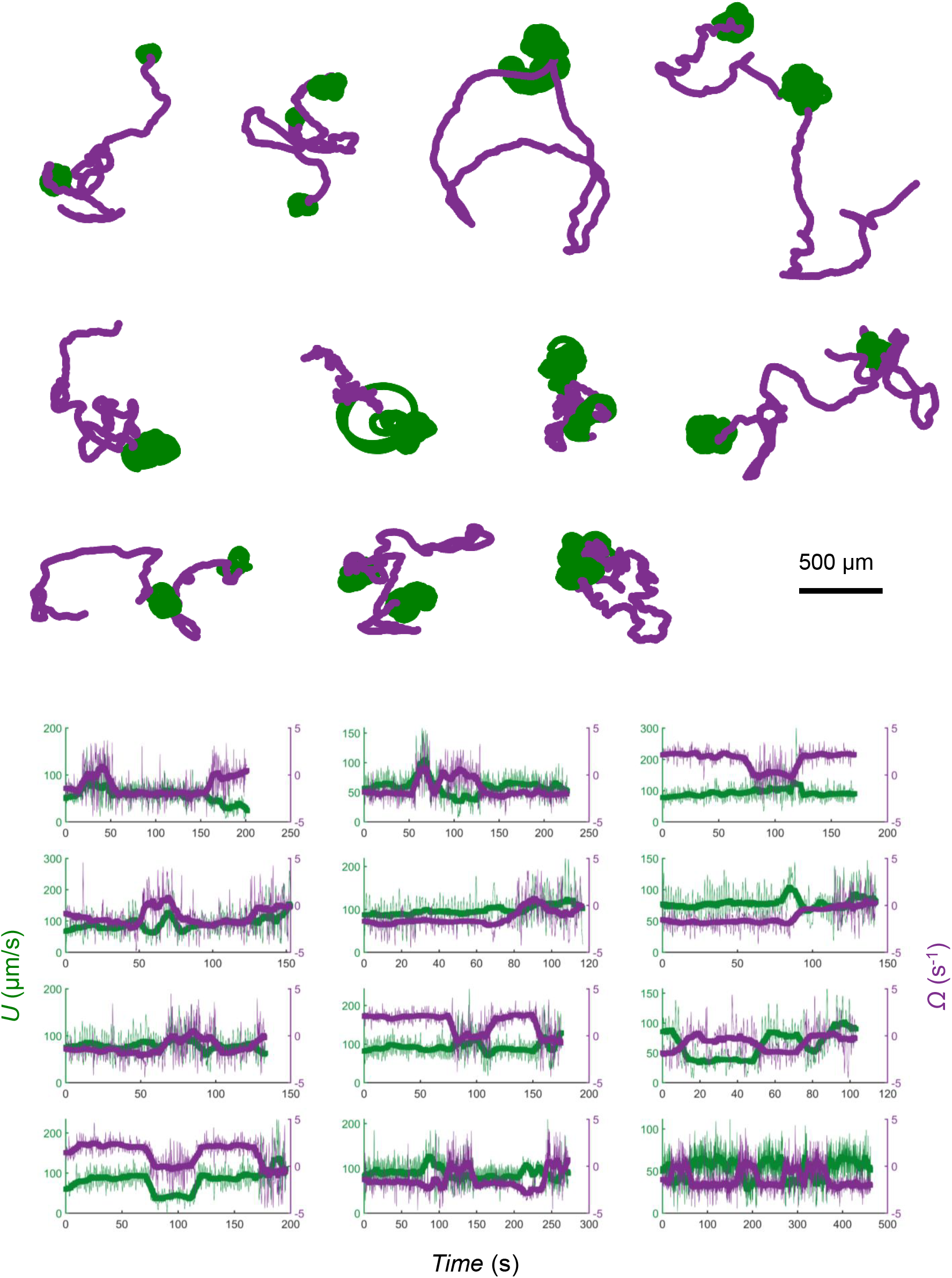
Trajectories of sperm circling and wandering. Circling and wandering behaviors with corresponding measurements of translational and rotational speeds. The transition between circling and wandering is observed in speed changes and rolling suppression.

**Figure S 3.**
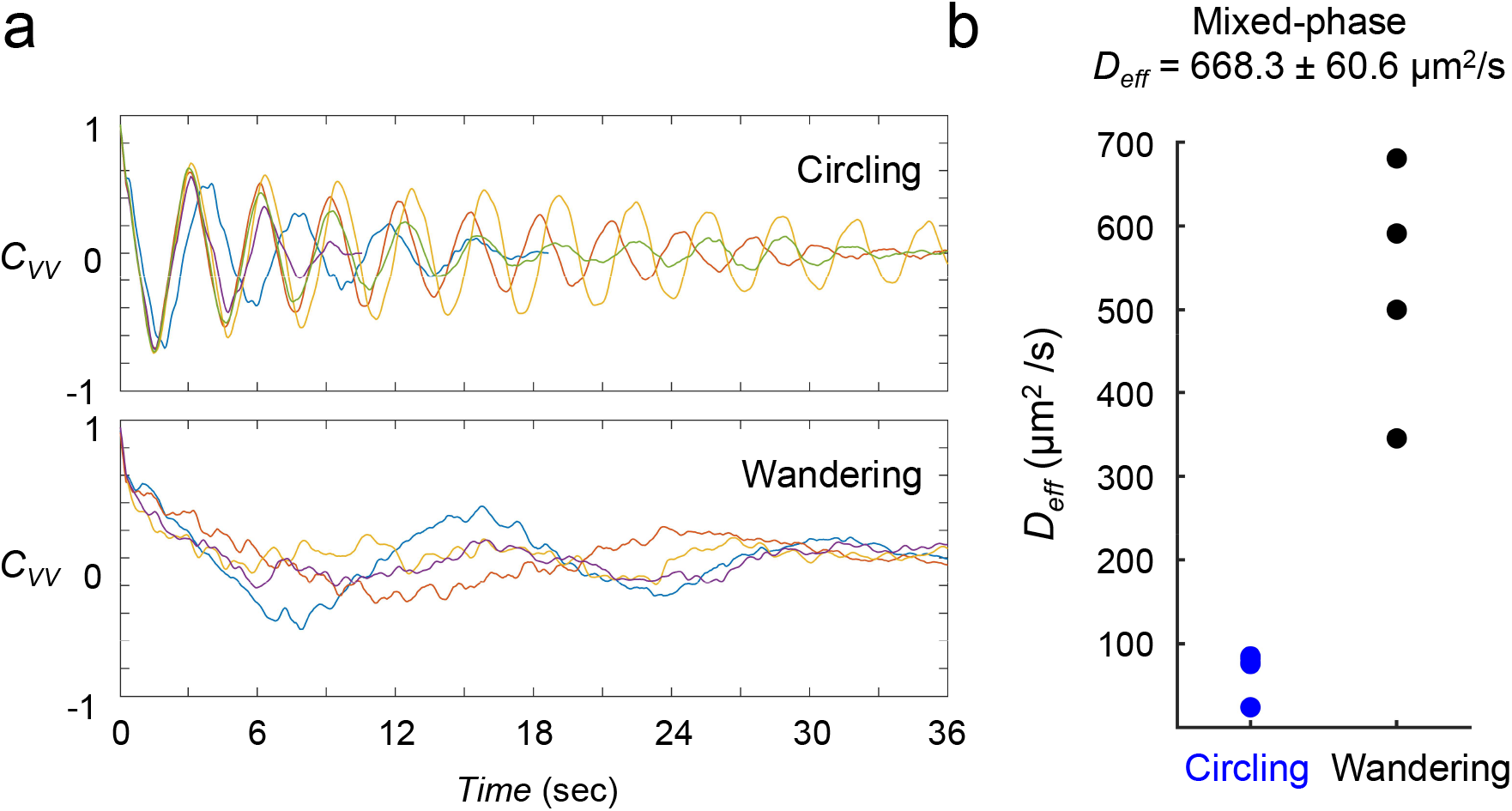
Constituent circling and wandering periods in mixed motility. (a) Velocity correlation function of constituent phases. Each color represents the trajectory of a constituent phase. (b) Effective diffusivities for each phase and the overall mixed phase.

**Figure S 4.**
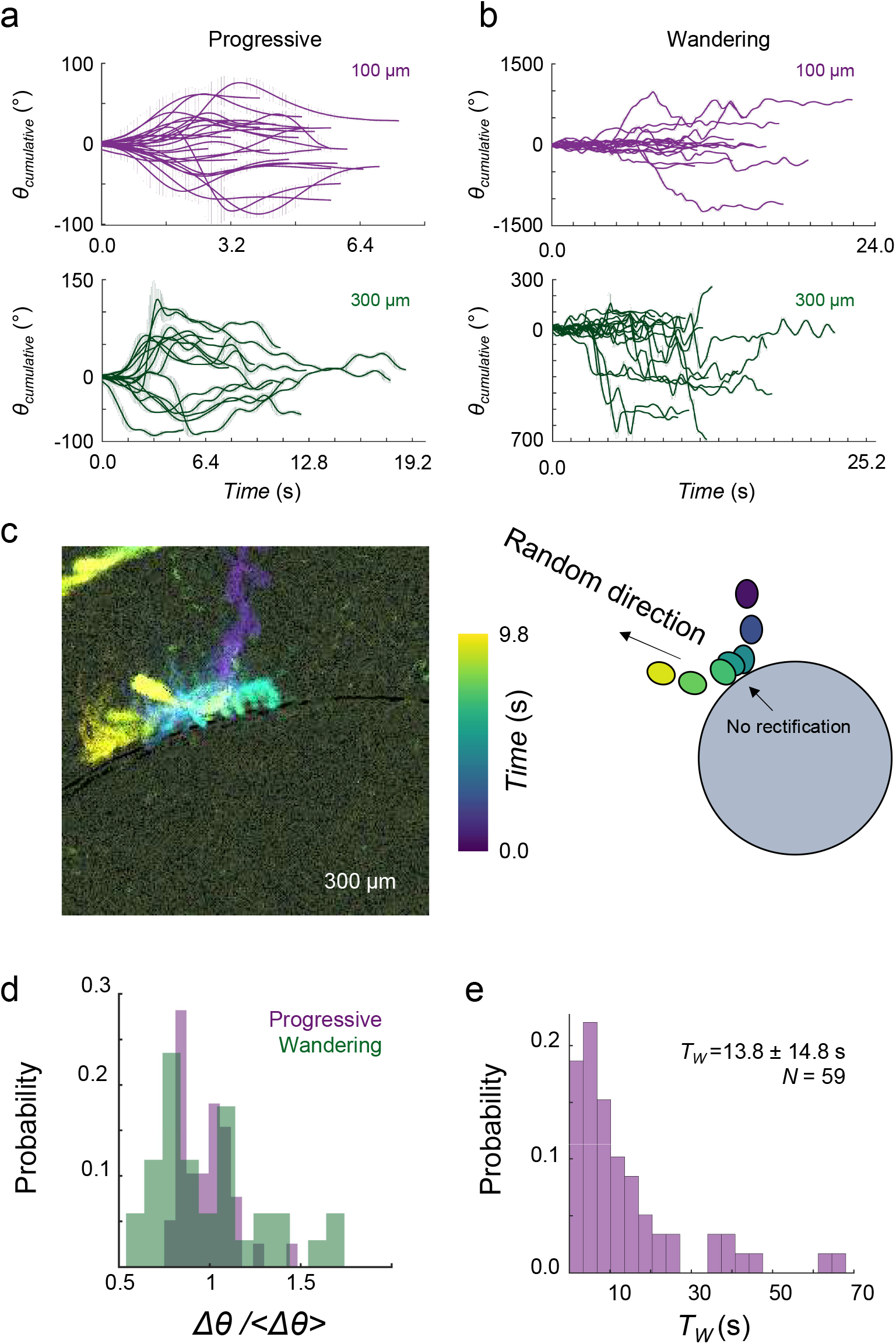
Wall interactions in the wandering phase. (a) Cumulative deflection angle (*θ*_cumulative_) for (a) progressive and (b) wandering sperm scattered from pillars of diameters *D* = 100 *µm* and *D* = 300 *µm*. (c) In the wandering phase, sperm contact length on the pillar is deterministically set by the deflection angle, while scattering occurs randomly without rectification by the pillar. (d) Histogram of sperm relative deflection angles in progressive and wandering phases. (e) Histogram of *T*_*w*_ for *N* = 59 trajectories.

**Figure S 5.**
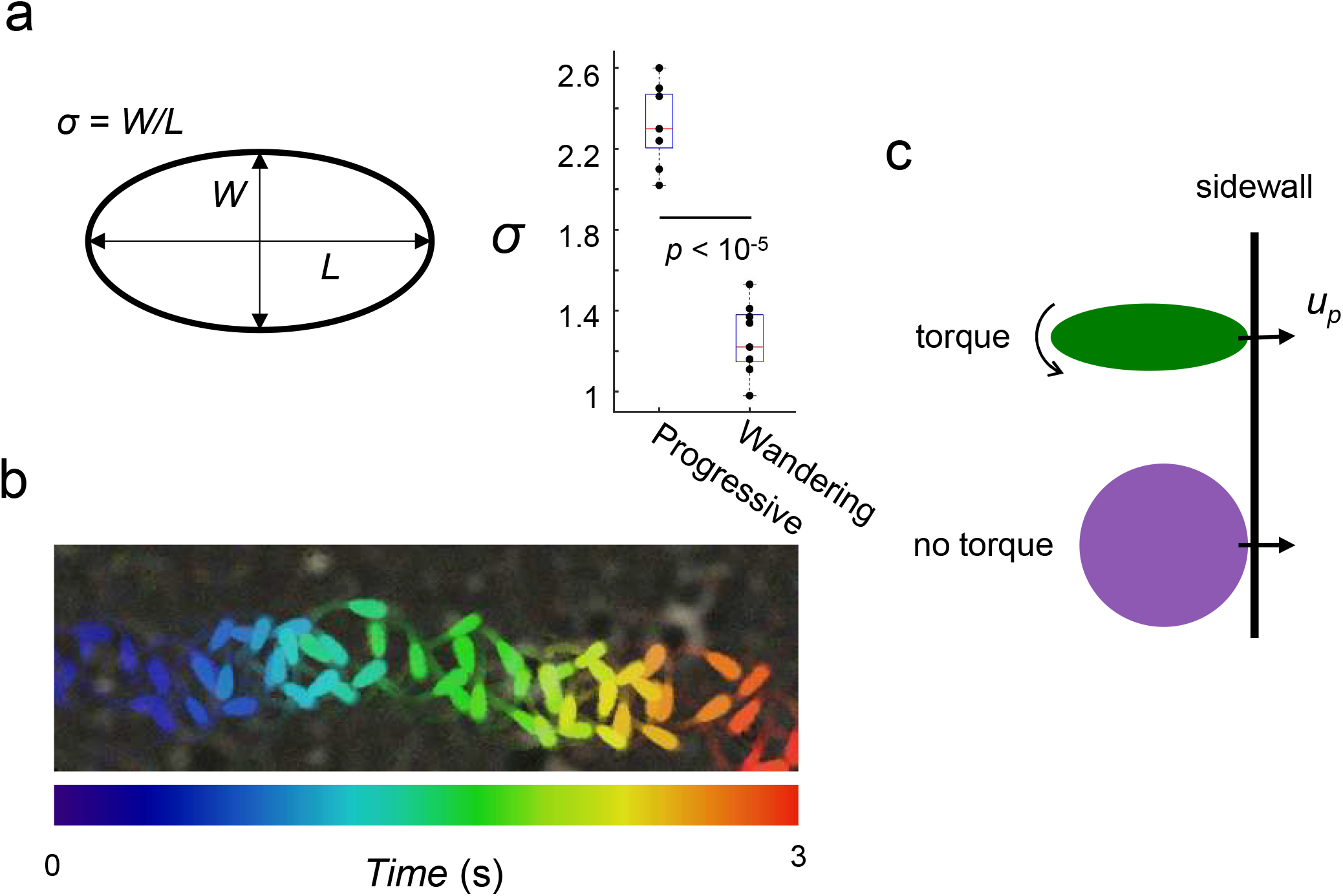
Sperm aspect ratio. (a) Aspect ratio measured from overlaid frames of sperm over a 600-millisecond period with 60-millisecond intervals, capturing both progressive and wandering phases. (b) Wandering sperm exhibit an effectively spherical shape compared to the elongated oval shape of progressive sperm. (c) High-aspect-ratio, oval-shaped swimmers experience an aligning torque upon collision with a physical boundary, whereas low-aspect-ratio, spherical swimmers experience significantly lower aligning torque.

**Figure S 6.**
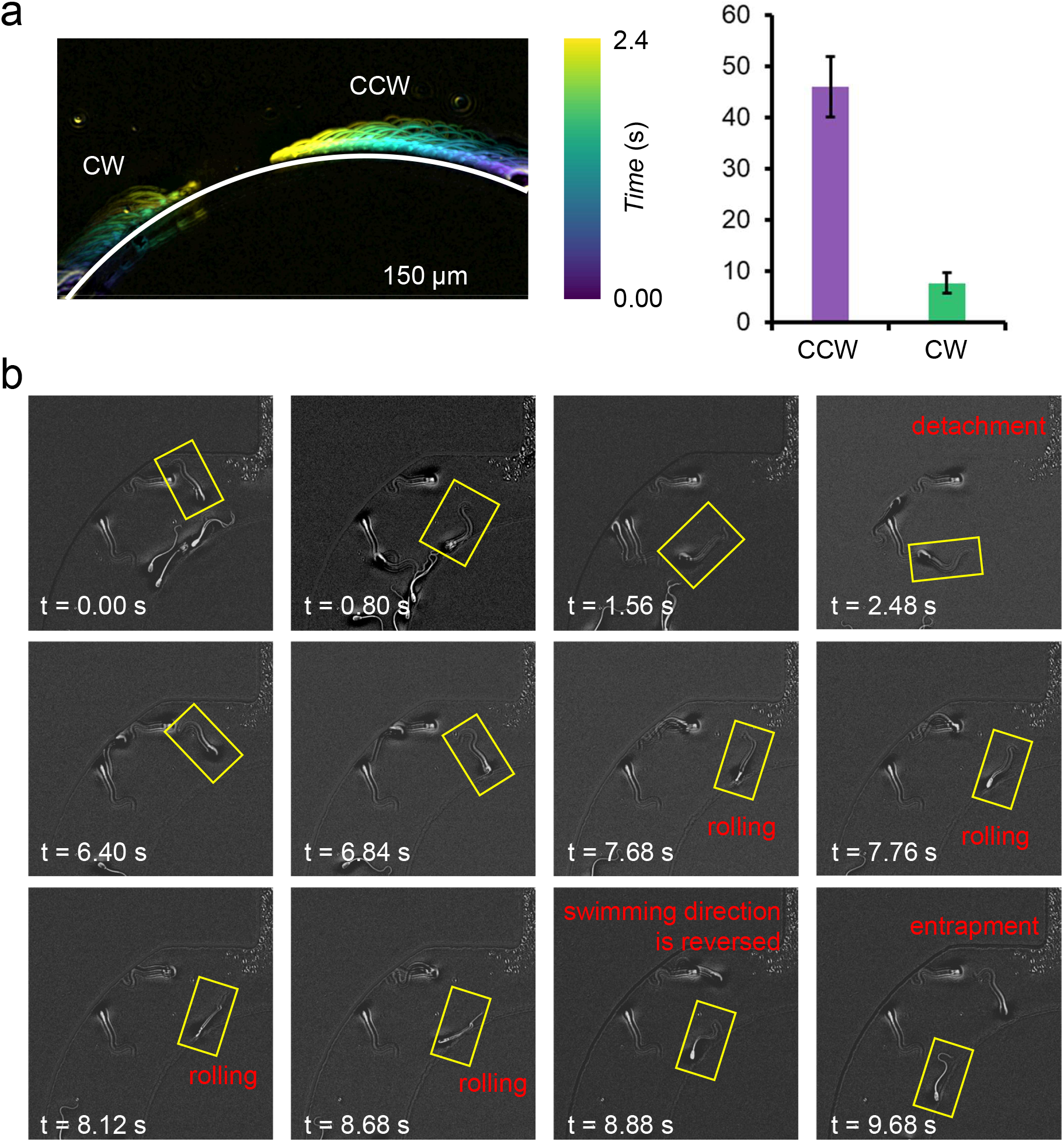
Characterization of Sperm Entrapment I. (a) Circling sperm with both clockwise (CW) and counterclockwise (CCW) chirality trapped around the *D* = 300 *µm* pillar. (b) Transitory rolling events reversed the chirality of sperm circling motion, leading to entrapment around the *D* = 300 *µm* pillar.

**Figure S 7.**
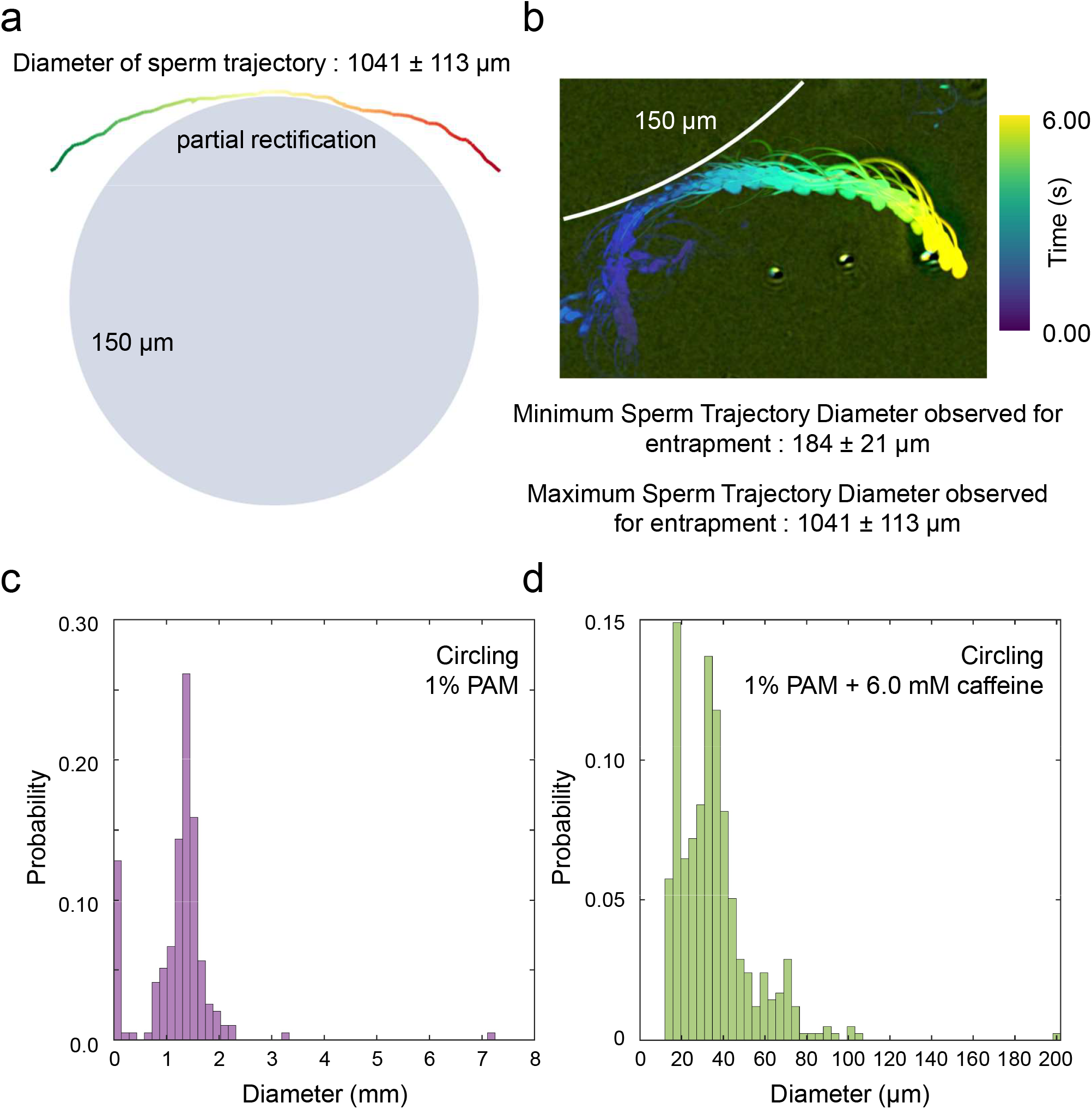
Characterization of Sperm Entrapment II. (a) Trajectory of a non-caffeine-treated circling sperm that is partially rectified by the pillar. (b) Trajectory of a non-caffeine-treated circling sperm that is not rectified by the pillar. Based on these measurements, we estimate the minimum and maximum trajectory diameters required for entrapment around the pillar. (c) Histogram of the sperm trajectory diameter in the circling phase without caffeine treatment. (d) Histogram of the sperm trajectory diameter in the circling phase with caffeine treatment (i.e., hyperactivation).

**Figure S 8.**
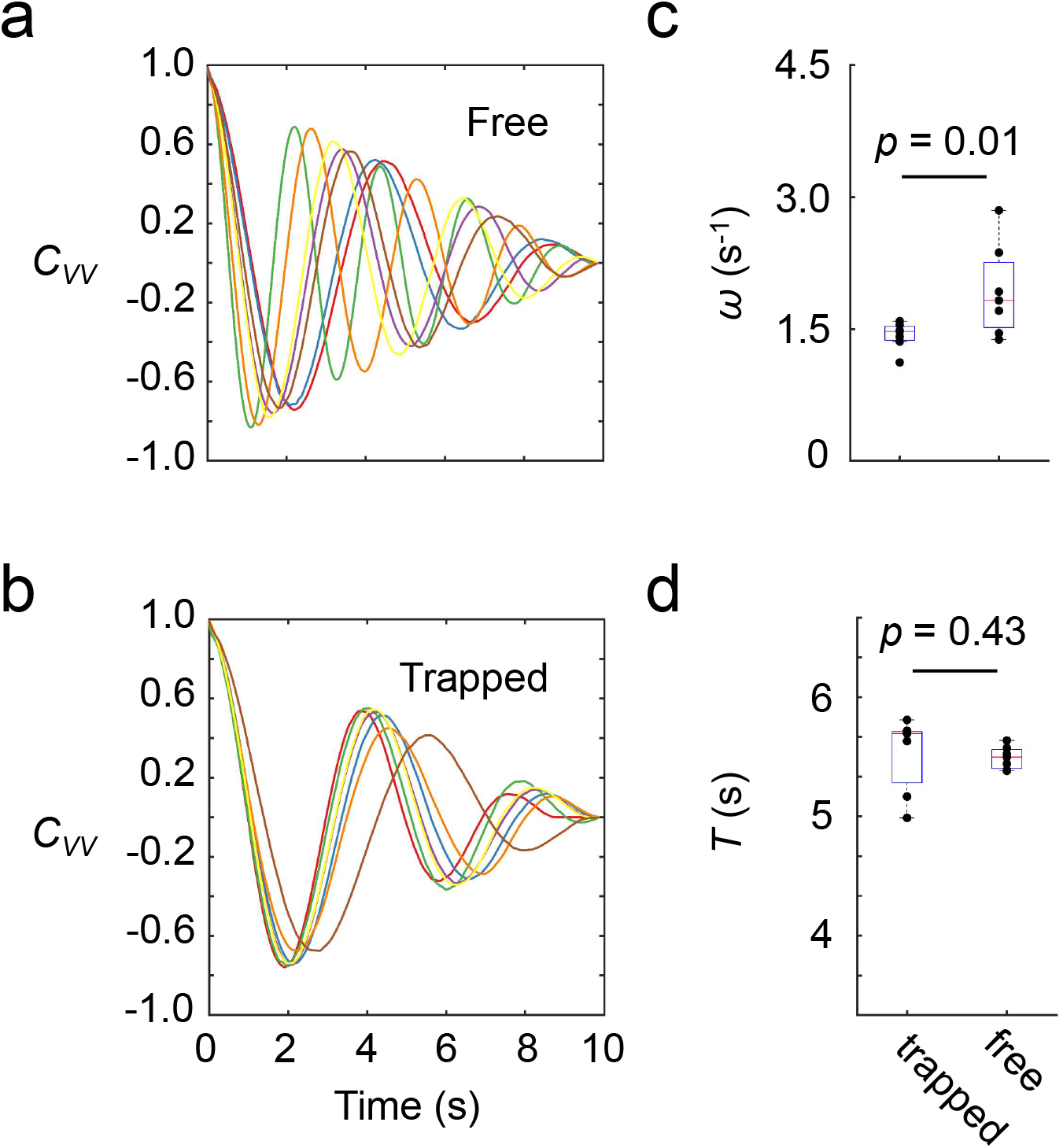
Characterization of Sperm Entrapment Around Small Pillars. (a) Velocity correlation function for individual caffeine-treated circling sperm before entrapment around the pillar. Each color represents an individual cell. (b) Velocity correlation function after entrapment. (c) Corresponding rotational speeds. (d) Corresponding relaxation times.

**Figure S 9.**
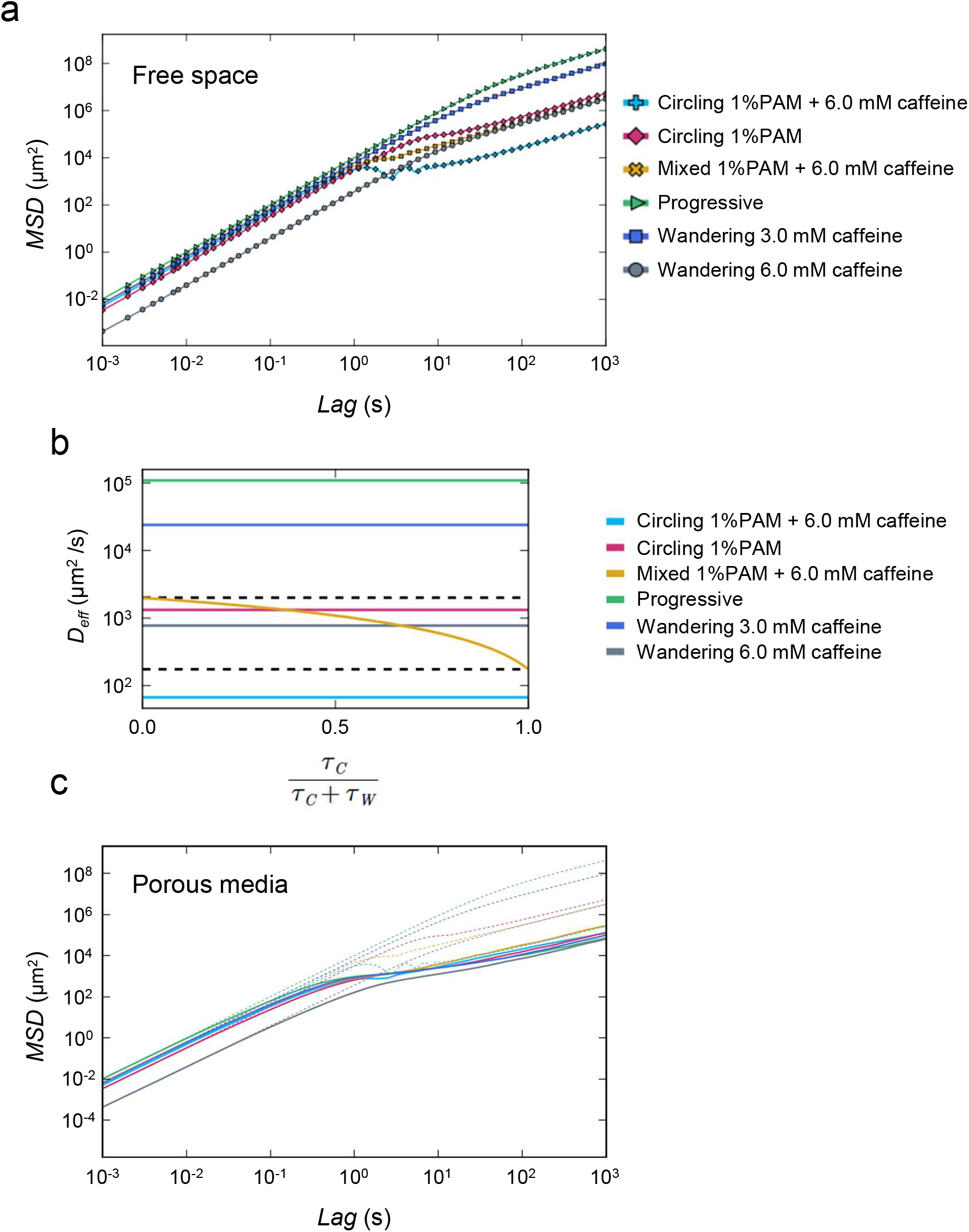
Mean-square displacements and effective diffusivity obtained from theory and simulations. (a) MSDs in free space computed for each motility phase. Lines correspond to the theoretical predictions and symbols to simulations. (b) Effective diffusivity *D*_eff_ of the mixed phase as a function of the fraction of time spent in the circling phase. (c) MSDs in porous media (solid lines) for *η* = 0.55 and mean chord length *l*_*h*_*/l*_*c*_ = 0.88, obtained from simulations. Dashed lines show the theoretical predictions for the MSD in free space for comparison.

